# Aging associated RNA loss impairs the dynamics of RNA and protein condensates

**DOI:** 10.64898/2026.02.16.705223

**Authors:** Anuva Rajappa, Twinkal Patel, Sumeru Sugata Raju, Nischitha Baliga, Rashmi Parihar, Ankita Chodankar, Akanksha Singh, Prarthana Ghosh Dastidar, K Venkatesan Iyer, Subramaniam Ganesh, Shovamayee Maharana

## Abstract

Altered biomolecular condensate dynamics are increasingly implicated in age-associated neurodegenerative disorders, yet the molecular principles governing these changes remain incompletely understood. Here, we demonstrate that aged cells across diverse cellular and organismal models harbor pre-formed, viscous, and persistent stress granules (SGs), which we term alt-SGs. These persistent condensates confer protection to senescent cells under fluctuating stress conditions. Comparative analyses reveal that alt-SGs exhibit elevated RNA-binding protein (RBP)-to-RNA ratios, underscoring a shift in condensate stoichiometry. By quantifying SG composition and dynamics in live cells and in vitro, we establish that RNA concentration is a key determinant of condensate material properties, molecular composition, and dissolution capacity. Importantly, aging cells display diminished absolute RNA concentrations due to reduced RNA metabolic activity. Reactivation of RNA metabolism restores RNA/RBP ratios within SGs and alleviates aging-associated phenotypes. Together, our findings highlight RNA metabolism as a central regulator of condensate stoichiometry and function, linking metabolic decline to altered phase behavior and cellular aging.

**Key findings:**

- Aging and senescence have Alt-SGs with increased protein/RNA Ratio
- Alt-SGs are persistent after stress removal in aging
- Aging is associated with reduced total cellular RNA
- Reactivating RNA metabolism rescues SG phenotypes
- Alt-SGs are protective for senescent cell

## Introduction

Aging at both the organismal and cellular levels is marked by a progressive decline in cellular and tissue function. At the molecular level, aging cells accumulate reactive oxygen species (ROS), misfolded proteins, and DNA and RNA damage (Chen et al. 2026; Nunomura et al. 2012). Concomitantly, aging is accompanied by profound structural and functional remodelling of cellular architecture. Key organelles undergo characteristic changes: mitochondria become energetically inefficient and more permeable, the Golgi apparatus fragments with impaired protein modification and vesicular trafficking, and endoplasmic reticulum mass and function decline.

Beyond membrane-bound organelles, biomolecular condensates constitute a critical and dynamic layer of cellular organisation that spatially and temporally coordinates biochemical reactions. Growing evidence indicates that these membraneless compartments undergo extensive remodeling during aging. RNA- and RNA-binding protein (RBP)–enriched condensates exhibit pronounced age-dependent changes, including nucleolar enlargement (Buchwalter and Hetzer 2017), altered nuclear speckle organization (Dion et al. 2025; Rhine et al. 2025), increased number and protein enrichment of P-bodies (Rieckher et al. 2018), and fusion of specialized RBP condensates into larger assemblies (Pushpalatha et al. 2022). Together, these observations identify condensate dysregulation as an emerging hallmark of cellular aging, although the molecular principles driving these alterations remain poorly understood.

RBPs that form condensates—including FUS, TDP-43, ataxin-2, FMRP, and hnRNPA1—also undergo age-associated transitions into solid-like cytoplasmic aggregates during neurodegeneration. Notably, many of these RBPs are core components of cytoplasmic RNA–RBP condensates known as stress granules (SGs). SGs assemble rapidly in response to diverse stresses, including oxidative stress, heat shock (Grousl et al. 2009; Hu et al. 2010; Huelgas-Morales et al. 2016), ER stress (Lyons et al. 2016), viral infection (Piotrowska et al. 2010), and nutrient deprivation (Reineke et al. 2018; T. Wang et al. 2022). Their nucleation is tightly coupled to stress-induced inhibition of translation initiation, most commonly mediated by phosphorylation of eIF2α (Kedersha et al. 2002), leading to ribosome dissociation and accumulation of untranslated mRNAs (Kedersha et al. 2002; Bounedjah et al. 2014; Van Treeck et al. 2018). These RNAs engage the SG scaffolds G3BP1/2, triggering multivalent interactions that drive condensate assembly.

SGs are thought to function as highly dynamic and reversible compartments that transiently sequester mRNAs and translation machinery and enable rapid recovery upon stress relief (Wheeler et al. 2016; Palangi et al. 2017; Hofmann et al. 2021). Consistent with this view, SG dynamics are progressively impaired with age: SGs exhibit reduced mobility in aged *C. elegans* (Lechler et al. 2017), and both SG assembly and disassembly are compromised during aging across systems (Omer et al. 2020; Dubinski et al. 2023). Despite these observations, **SG formation, dissolution, and material properties have not been quantitatively examined in aging cells in real time**, leaving unresolved whether SGs serve as precursors to pathological RBP aggregation or represent a parallel, independent state.

Here, using cellular, organismal, and in vitro reconstitution models, we show that aging fundamentally alters SG composition and biophysical properties. Across systems, aged cells form SGs enriched in RBPs but depleted of RNA, resulting in increased condensate viscosity and persistence. Strikingly, we demonstrate that **the RNA-to-RBP ratio within condensates is a key determinant of their material state**, and that restoring RNA levels reverses aberrant SG dynamics and alleviates cellular aging phenotypes. **This work provides the first direct evidence that age-associated changes in RNA–RBP stoichiometry actively govern condensate dynamics, uncovering RNA homeostasis as a central regulator of biomolecular condensates during aging.**

## Results

### Senescent cells are prone to form persistent and viscous SGs

SGs are composed of RNA, scaffold RBPs, and phase separation–prone proteins such as hnRNPA1, FUS, and TDP-43, which are also implicated in age-related neurodegenerative diseases. To compare the dynamics of SG formation between young and aged cells, we induced senescence in HeLa cells that stably express mCherry-tagged G3BP1, a key SG scaffold protein (Poser et al. 2008; Guillén-Boixet et al. 2020). Senescence in HeLa cells was triggered by BrdU-induced DNA damage (Methods; Supp Fig. 1A). As expected, BrdU-treated cells displayed all previously reported senescence markers (Ross et al. 2008; Nair et al. 2015), including enlarged nuclei, increased cell size, elevated senescence-associated β-galactosidase activity, and higher expression of established senescence markers (Supp Fig. 1A).

To quantify SG formation dynamics, we imaged the G3BP1-mCherry expressing HeLa cells during the exposure to sodium arsenite (NaAsO₂) (Guillén-Boixet et al. 2020) (Fig. 1A,B). Remarkably, even in the absence of oxidative stress, approximately 40% of senescent HeLa cells contained pre-formed SGs (Supp Fig. 1C). These pre-formed SGs in senescent cells were typically ∼0.200 μm² in area or ∼0.5 μm in diameter (Supp Fig. 1C). Based on this observation, we hypothesize that senescent cells are inherently more prone to SG formation. To compare SG formation rates, we initially imaged stressed cells every minute at 37 °C, which led to rapid cell shrinkage (Supp. Fig. 1B), particularly in senescent cells. To minimize imaging-induced stress, subsequent experiments were performed at 25 °C, slowing SG formation and reducing cell shrinkage. Control cells did not contain pre-formed SGs and began to display visible SGs after 25 minutes. By 80 minutes, nearly all control cells had formed SGs (Fig. 1C). SG formation was further quantified by normalizing the SG area fraction per cell (total SG area relative to cytoplasmic area) to the initial time point. The analysis showed that SG area in control cells increased approximately two-fold compared to the baseline, when SGs were absent (Fig. 1D). Likewise, G3BP1 enrichment within SGs (ratio of G3BP1 concentration in SGs to that in the cytoplasm) reached its maximum at 80 minutes (Fig. 1E). However, prolonged stress beyond 80 minutes resulted in a sharp decline in the proportion of SG-positive cells (Fig. 1C) and a reduction in SG area per cell (Fig. 1D), while G3BP1 enrichment decreased only slightly, consistent with previous reports (T. Wang et al. 2022; Adachi et al. 2024).

**Figure 1:**
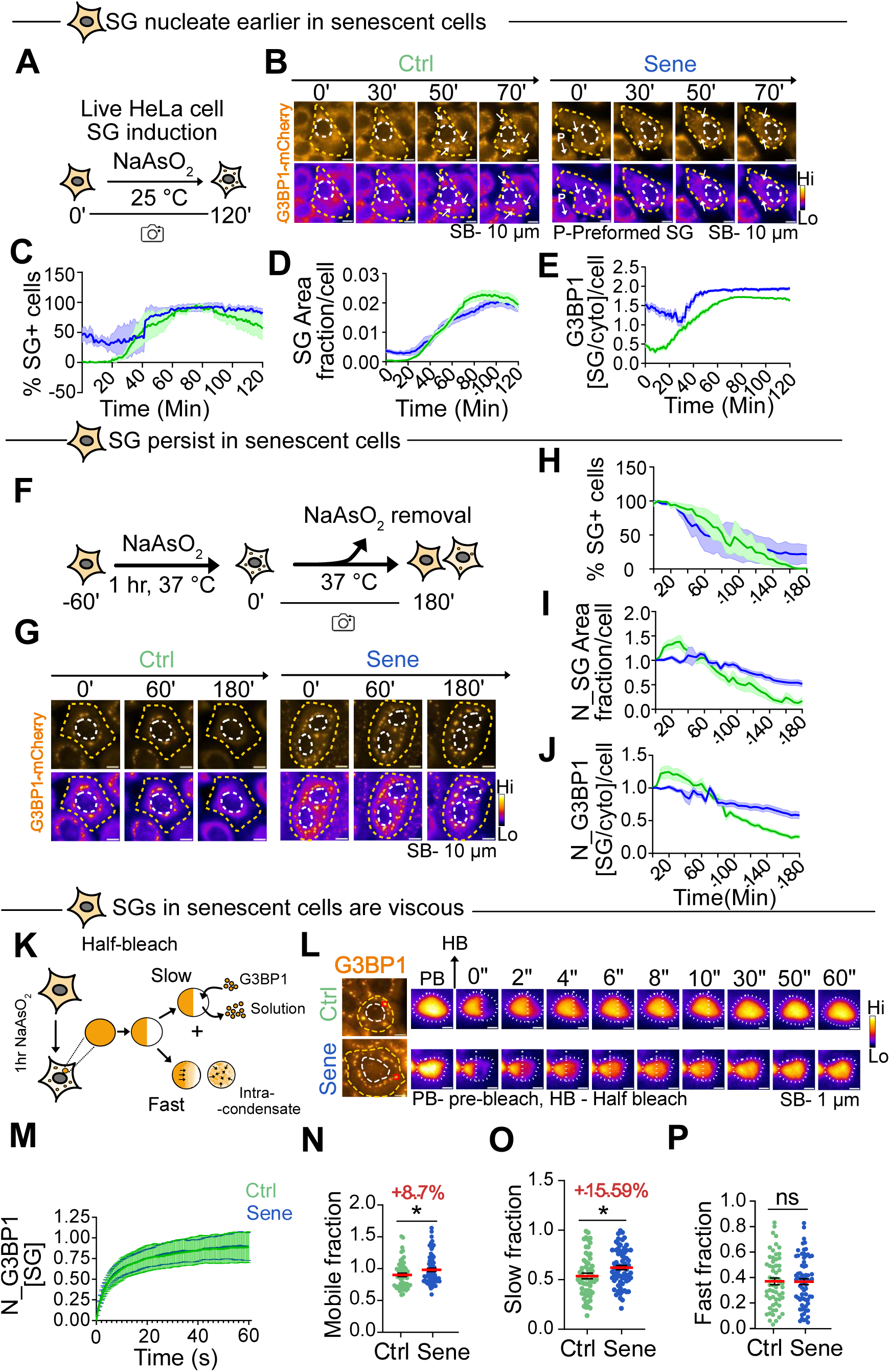
Senescent cells are prone to form persistent and viscous SGs. (A) HeLa cells expressing G3BP1-mCherry were imaged during treatment with 0.25 mM NaAsO_2_ for 120 mins at 25 °C to track SG formation. (B) Montage of control and senescent HeLa cells expressing G3BP1-mCherry under oxidative stress. Images at 0′ are before NaAsO₂; subsequent frames show stress-induced G3BP1 foci (white arrows). Quantification of (C) percentage of cells containing SGs, (D) SG area fraction, and (E) mean G3BP1 enrichment within SGs per control and senescent cell during stress exposure. For each condition, ≥200 cells were analyzed across 20 independent time-lapse movies; data represent mean ± SEM. (F) HeLa cells expressing G3BP1-mCherry were stressed with NaAsO₂ (60 min, 37 °C) to induce SGs; following stress removal, SG dissolution was monitored by live-cell imaging for 180 min. (G) Montage of SGs in G3BP1-mCherry expressing control and senescent HeLa cells following NaAsO₂ removal. 0’ shows SGs formed after 1 h of stress, prior to stress withdrawal; subsequent panels depict SG dissolution during the indicated time points. Quantification of (H) percentage of cells containing SGs, (I) SG area fraction, and (J) mean G3BP1 enrichment within SGs per control and senescent cell during stress recovery. For each condition, ≥200 cells were analyzed across atleast 15 independent time-lapse movies; data represent mean ± SEM. (K) Half-FRAP of SGs tagged with G3BP1-mCherry, formed after 1 h of NaAsO₂ treatment at 37 °C. The schematic illustrates two mechanisms of fluorescence recovery: (i) slow diffusion of SG components from the cytosol into the condensate and (ii) rapid molecular rearrangement within the condensate. (L) Montage of half-frapped G3BP1-mCherry–labeled SG induced by 1 hour of NaAsO₂ stress used for calculation of mean recovery curve. Images show SGs before bleaching (PB) and after a half-bleach (HB). Selected granules were half-bleached for 50 milliseconds, and fluorescence recovery was recorded for 60 seconds. (M) Mean FRAP recovery curves plotted by quantifying normalized mean G3BP1 intensity in half-bleached SGs (mean ± SD; ≥60 curves per condition). (N) Mobile fraction of SGs derived from individual recovery curves. Quantification of (O) slow fraction (diffusion from solution) and (P) fast fraction (intracondensate rearrangement) contributing to overall mobility. Statistical significance determined by Mann–Whitney U test (*P < 0.05, **P < 0.01, ***P < 0.001).

Senescent cells exhibited markedly different SG formation dynamics. Nearly half (∼48%) of senescent cells contained pre-formed SGs, and by 45 minutes, almost all senescent cells had developed SGs (Fig. 1C). Despite the presence of pre-formed SGs, the increase in SG area fraction over time was less pronounced in senescent cells compared to controls (Fig. 1D). Both control and senescent cells reached their respective peaks of SG formation at 80 minutes; however, senescent cells had a lower amount of total SGs than in controls, consistent with previous findings (Dubinski et al. 2023). Notably, G3BP1 enrichment within SGs peaked earlier in senescent cells than in controls, likely reflecting the contribution of pre-formed SGs (Fig. 1E).

Unlike in control cells, prolonged stress after 80 minutes in senescent cells did not decrease the number of SG-positive cells (Fig. 1C). Almost all senescent cells retained SGs even at 120 min. The normalized area fraction (Fig. 1D) and G3BP1 SG enrichment (Fig. 1E) also remained high during the prolonged stress from 80-120 minutes. SGs are liquid-like condensates that grow in size through the diffusion of RNAs and RBPs from solution to condensates, as well as through the fusion of condensates. The fewer SG fusions in senescent cells might also be the cause of smaller SGs as quantified by the frequency distribution of the SG sizes in control and senescent cells (Supp Fig. 1E). Additionally, the fusion of condensates may also reveal their liquid-like properties. We found fewer SG fusion events in senescent cells than in control cells when quantifying fusion events in equal numbers of cells over 120 minutes of imaging (Supp Fig. 1D).

Cells often experience stress and form SG, which sequesters the translating mRNA (Glauninger, Bard, Wong Hickernell, et al. 2025), but these SG resolve after the removal of the stress conditions (Wheeler et al. 2016; Palangi et al. 2017; Hofmann et al. 2021), restoring translation to physiological levels (Das et al. 2022; Sugawara et al. 2024). The presence of pre-formed SG suggested a defective SG resolution after stress removal. To measure SG dissolution, we induced SGs with an hour of sodium arsenate treatment and then monitored their disassembly for the next 180 minutes after removal of stress (Fig. 1F,G). In control cells, the percentage of SG-positive cells began to decline after approximately 35 minutes of stress removal, and by 160 minutes, almost no cells had SGs. In senescent cells, SG dissolution was initially rapid, with a marked decrease in SG-positive cells within the first 60 minutes; however, the process then slowed, and even after 180 minutes, ∼30% of senescent cells still retained SGs. Consistent with this, the normalized total SG area fraction and the normalized SG enrichment of G3BP1 per cell in control conditions began to decline sharply after 40 minutes and maintained a downward trend (Fig. 1H, I). At 180 min post-stress removal, senescent cells had 40% higher SG area fraction and 30% higher G3BP1 enrichment in the SG as compared to SG in control cells (Fig. 1J). Since live cell imaging of SG dissolution causes additional stress due to exposure to light and requires fluorescently tagged G3BP1, we also measured the dissolution of endogenous SGs in HeLa cells at 1.5, 3 and 4 hours of recovery from an hour of stress (Supp Fig. 1F, G). Almost 54.2% of the senescent cells had SG after 4 hours of stress removal, as compared to only 29.5% of cells that were left with SG in control cells (Supp Fig. 1H). Hence, live-cell imaging and quantification of SG formation and dissolution revealed that senescent cells were more prone to form SGs, had pre-formed SGs, and exhibited defective SG dissolution mechanisms, leading to the persistence of SGs. As the SGs in senescent cells were highly enriched for SG scaffold proteins, smaller, and persistent, to distinguish them from SG in control cells, we will hereafter refer to them as **alt-SGs**.

A large fraction of SGs resolve by reversal of phase separation after stress removal as the SG-nucleating RNA returns to translating ribosomes, and remaining SGs are cleared by active processes like autophagy (Das et al. 2022; Sugawara et al. 2024; Buchan et al. 2013). Material property of the SGs has been speculated to control their fate on stress recovery, although no quantitative measurements have been taken till now. We postulated that Alt-SGs may have altered material properties, which can make the reversal of phase separation difficult. To assess SG material properties, we performed half-FRAP (fluorescence recovery after photobleaching) on SGs larger than 0.25 μm in diameter in both control and senescent cells. Recovery of half-FRAPed condensates occurs through two processes: (a) rapid intra-condensate mixing (fast fraction), and (b) slower diffusion of fluorescent proteins from the surrounding cytoplasm into the condensate phase (slow fraction) (Fig. 1K). The Half FRAP recovery curves of SGs in control and senescent cells appeared very similar (Fig. 1L, M; Supp 1I, J). To quantify the fast and slow fractions, we next fitted the Half FRAP recovery curves with a double exponential function. The total mobile fraction was about 8.7% higher for Alt-SGs than the SGs (Fig. 1P). The contribution of the slow fraction to the total mobile fraction was ∼15.59% higher in the case of Alt-SGs from senescent cells compared to SGs. The time scale of both fast and slow fractions was very similar for the control and senescent cells (Supp 1K, L). The increased contribution of the slow fraction suggested that most recovery occurred through diffusion from the cytoplasm to the condensate rather than faster intra-condensate mixing (Fig. 1N, O), suggesting a more viscous nature of the condensate.

Our studies have for the first time found that the material property of the SGs are connected to their dissolution dynamics in stress recovery. And senescent cells have more enriched and viscous alt-SGs, which make them persistent in nature.

### Aged cells across different model organisms have enriched SGs

We confirmed the enriched and persistent Alt-SGs observed in live cells by examining endogenous SGs through immunofluorescence. G3BP1 was highly enriched in senescent cells at 15, 30, 60, and 120 minutes (Fig. 2A, B; Supp. Fig. 2A-C). The similarity in SG phenotypes between fixed and live cells suggests that endogenous SGs in senescent cells are also Alt-SGs. Notably, the SG area fraction normalized to cell area was higher in fixed cells compared to live cells (Fig. 2C). However, fixed-cell analysis did not detect pre-formed SGs, likely due to fixation, permeabilization, and washing steps that can remove smaller SGs during immunofluorescence.

**Figure 2:**
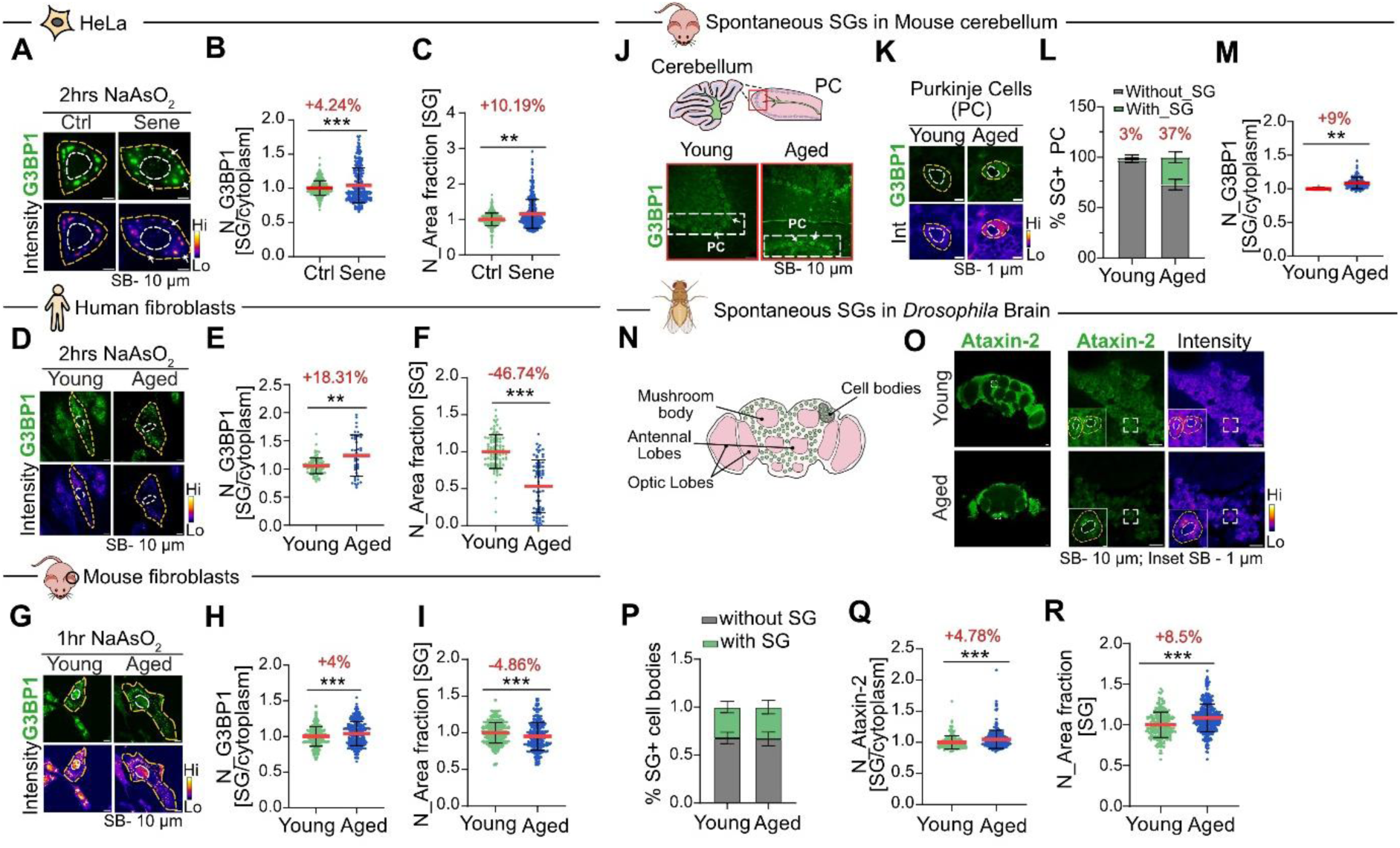
Aged cells across different model organisms have enriched SGs. (A) SGs in control and senescent HeLa cells after 2 h of 0.25 mM NaAsO₂ treatment at 37 °C, detected with antibodies against endogenous G3BP1. White arrows highlight smaller SGs observed in senescent cells. Quantification of (B) G3BP1 enrichment within SGs per cell and (C) SG area fraction per cell. For each condition, >200 cells were analyzed; data represent mean ± SD. (D) SGs in young and aged human fibroblasts after 2 h of 0.25 mM NaAsO₂ treatment at 37 °C, detected with antibodies against endogenous G3BP1. Quantification of (E) G3BP1 enrichment within SGs per fibroblasts (F) SG area fraction. For each condition, at least 50 cells were analyzed; data represent mean ± SD. (G) SGs in young and aged mouse fibroblasts after 1 h of 0.25 mM NaAsO₂ treatment at 37 °C, detected with antibodies against endogenous G3BP1. Quantification of (H) G3BP1 enrichment within SGs per mouse fibroblasts (I) SG area fraction. For each condition, >200 cells were analyzed; data represent mean ± SD. (J) Detection of spontaneous SGs in cerebellar sections from 2-month-old (young) and 18-month-old (old) mice using G3BP1 staining. The schematic shows the granular layer containing the Purkinje cell (PC) layer (black dotted line); the red box highlights a magnified view of this region. White dotted boxes and arrows indicate PCs in young and aged cerebellar sections. (K) Spontaneous SGs in PCs from young and aged mouse brain sections stained for endogenous G3BP1. (L) Bar graph showing the percentage of SG-positive PCs. (M) Quantification of G3BP1 enrichment in spontaneous SGs per PC in young and aged cerebellum. For each condition, ≥30 PCs (aged) and ≥10 PCs (young) were analyzed cumulatively from at least three mouse brains; data represent mean ± SEM. (N) Schematic showing neuronal cell bodies adjacent to the Mushroom Bodies and Antennal Lobes. Young (3-day-old) and aged (40-day-old) flies were analyzed for spontaneous SGs in the neuronal cell bodies. (O) Spontaneous SGs in neuronal cell bodies from young and aged mouse brain sections stained for endogenous Ataxin-2; white dotted box highlights cell bodies containing SGs. (P) Bar graph showing the percentage of SG-positive neuronal cell bodies. (Q) Quantification of Ataxin-2 enrichment in spontaneous SGs per neuronal cell body, (R) SG area fraction in young and aged Drosophila. For each condition, >100 neuronal cell bodies were analyzed cumulatively from at least 18 Drosophila brains. All data represent mean ± SD unless specified and are normalised to the mean intensity values of respective controls. Statistical significance determined by Mann–Whitney U test (*P < 0.05, **P < 0.01, ***P < 0.001).

Next, we investigated the SG properties in naturally aged cells, i.e., primary human and mouse fibroblasts derived from young and aged individuals. We used human foreskin primary fibroblasts isolated from young (11-year-old) and aged donors (75-year-old) (Roy et al. 2020), as well as primary fibroblasts isolated from the ears of young (2-month-old) and aged (18-month-old) mice. SG enrichment of G3BP1 was higher in both human and mouse organismal aging models (Fig. 2D, E and G, H). The total SG area fraction per cell was now lower in both aged human and mouse fibroblasts when compared to the younger fibroblast (Fig. 2F, I). We also compared the size distribution of the endogenous SGs and Alt-SGs in human, mouse fibroblast and HeLa cells. In both mouse and human fibroblasts, the endogenous Alt-SGs were smaller (Supp Fig. 2E, F; Supp Fig. 2G); however, in HeLa cells, they were larger (Supp Fig. 2D). These quantifications suggested that SG enrichment is a more consistent phenotype across different organisms and aging models, whereas other SG properties, such as per-cell SG area fraction and SG size, are variable phenotypes. We next probed the SG properties in the Purkinje cells (PC) of the cerebellum (Fig. 2J, K) from young and aged mice which were formed spontaneously without application of additional stress. We found almost 37% of PCs in old mice had SGs as opposed to only 3% of PCs with spontaneous SGs (Fig. 2L). The SGs in older mice were also about 9% more enriched for G3BP1 enrichment as compared to spontaneous SGs in younger mice (Fig. 2M), concretizing our results of enriched SGs as a more consistent phenomenon in mammals.

Since SGs and their scaffold proteins are conserved across eukaryotes (Yao et al. 2024), we examined whether SG phenotypes change with age in *Drosophila*. Brains from young (3-day) and aged (40-day) flies were stained for Ataxin-2, a key SG protein (Bakthavachalu et al. 2018; Singh et al. 2021) (Fig. 2N). Spontaneous Ataxin-2–enriched SGs were detected in neuronal cell bodies (Fig. 2O). Unlike mouse PCs, the frequency of SG-positive cell bodies was similar in young and aged flies (Fig. 2P). However, Ataxin-2 enrichment was higher in aged brains (Fig. 2Q), consistent with mammalian cells (Fig. 2B, E, H). The area fraction of the spontaneous SGs was higher in the aged (Fig. 2R) cell bodies. Similar trends were observed in principal cells of Malpighian tubules (MT), where aged flies showed more enriched spontaneous SGs than young ones (Supp. Fig. 2H-J). Our data across organisms, tissues, and aging models show that Alt-SGs, which are enriched for scaffold RBPs, are viscous and persistent, which also makes aged tissues more prone to form SGs.

### Alt-SGs have more prion-like RBPs and less RNA

Till now, we found that Alt-SGs are enriched in G3BP1, a key scaffold protein required for SG nucleation (Yang et al. 2020; Guillén-Boixet et al. 2020). G3BP1 drives SG assembly only in the presence of free RNA (Kedersha et al. 2002; Bounedjah et al. 2014; Van Treeck et al. 2018). Other SG-associated RBPs (Gilks et al. 2004; Aulas et al. 2012; Shelkovnikova et al. 2013) also contribute to SG formation. Based on these findings, we next examined whether RNA concentration and the abundance of SG-associated RBPs differ in Alt-SGs compared with SGs in young cells. We quantified polyA mRNA within SGs using fluorescence in situ hybridization (FISH) with oligo-dT probes. Alt-SGs in senescent HeLa cells showed a ∼20% reduction in polyA enrichment (Fig. 3A, B), while human fibroblasts displayed only a marginal decrease compared to control SGs (Supp Fig. 3Q).

**Figure 3:**
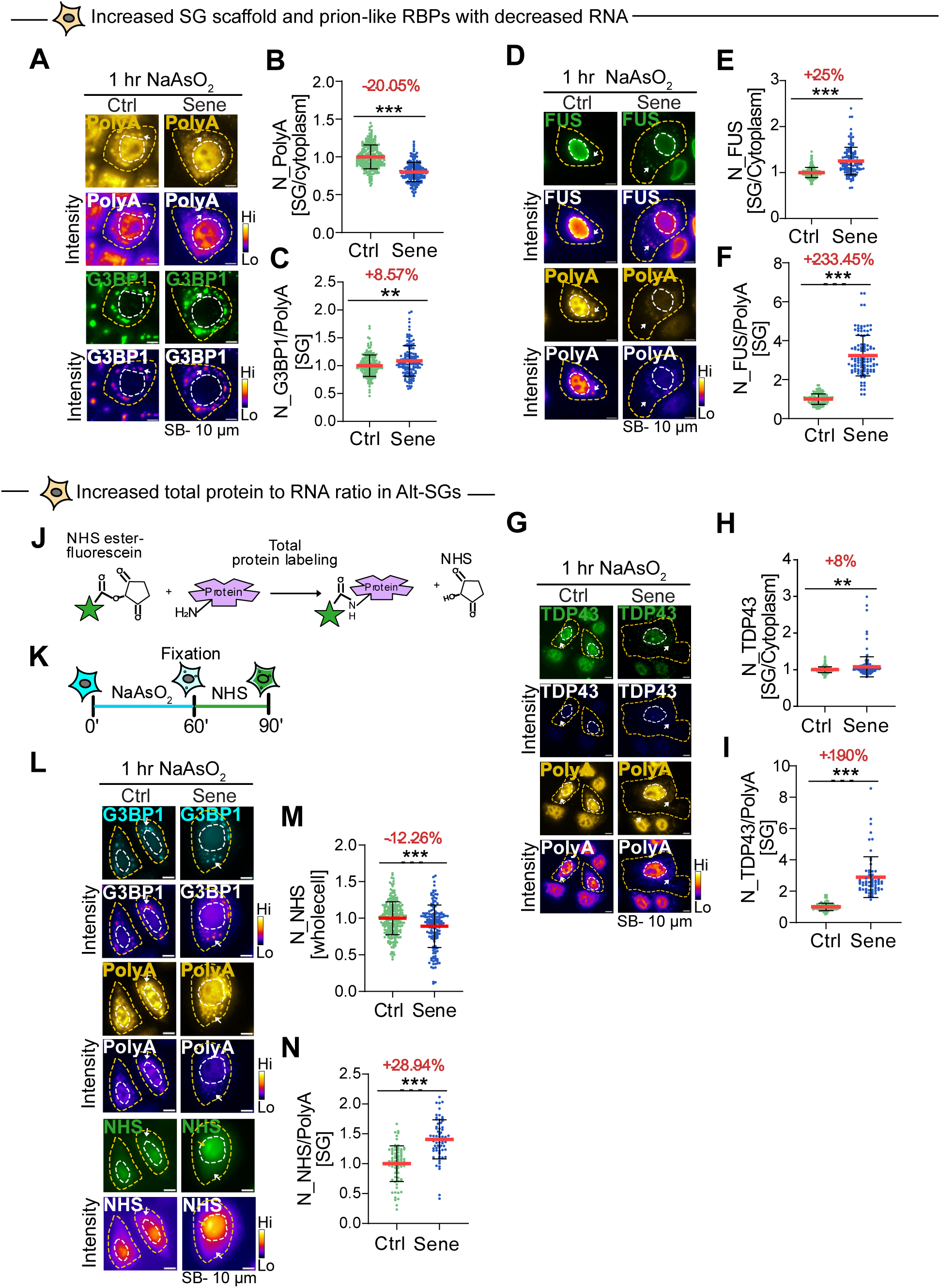
Alt-SGs have more prion-like RBPs and less RNA. (A) Detection of G3BP1 and polyA-tailed mRNA using fluorescent oligo(dT) FISH after 1 h of 0.25 mM NaAsO₂ stress in HeLa cells. (B) Quantification of polyA enrichment in SGs, expressed as the ratio of mean polyA intensity in SGs to cytoplasm. (C) Ratio of G3BP1 to polyA within SGs, calculated as mean G3BP1 intensity divided by mean polyA intensity per cell. For each condition, >170 cells were analyzed. (D) Detection of FUS and polyA-tailed mRNA after 1 h of 0.25 mM NaAsO₂ stress in HeLa cells. (E) Quantification of FUS enrichment in SGs, expressed as the ratio of mean FUS intensity in SGs to cytoplasm, and (F) the ratio of FUS to polyA within SGs, calculated per cell. For each condition, >100 cells were analyzed. (G) Detection of TDP43 and polyA-tailed mRNA after 1 h of 0.25 mM NaAsO₂ stress in HeLa cells. (H) Quantification of TDP43 enrichment in SGs, expressed as the ratio of mean TDP43 intensity in SGs to cytoplasm, and (I) the ratio of TDP43 to polyA within SGs, calculated per cell. For each condition, at least 100 cells were analyzed. (J) Schematic illustrating protein labeling by NHS ester–fluorescein, which covalently reacts with primary amines at peptide N-termini or lysine side chains to attach fluorescein. (K) Schematic of HeLa cells subjected to 30 min NHS treatment following 1 h of 0.25 mM NaAsO₂ stress, prior to fixation. (L) HeLa cells stained for G3BP1, polyA RNA, and total protein by NHS-fluorescein labeling. (M) Quantification of total protein, expressed as mean NHS-fluorescein intensity per cell. (N) Ratio of total protein to polyA within SGs, calculated as mean NHS-fluorescein intensity divided by mean polyA intensity per cell. For each condition, ≥155 cells were analyzed. All data represent mean ± SD unless specified and are normalized to mean intensity values of respective conditions. Statistical significance determined by Mann–Whitney U test (*P < 0.05, **P < 0.01, ***P < 0.001).

Previous work, including ours, has established that low RNA concentration promotes condensate nucleation (Maharana et al. 2018; Banerjee et al. 2017; Tejedor et al. 2021; de Vries et al., 2024) and prevents their transition into solid-like states (Maharana et al. 2018; Tejedor et al. 2021; Kim et al. 2023). To explain the viscous and persistent Alt-SGs described earlier, we next examined the RBP-to-RNA ratio within SGs. In Alt-SGs, the G3BP1-to-polyA ratio was elevated in both senescent HeLa cells (Fig. 3C) and aged human fibroblasts (Supp Fig. 3Q). This increase is not due to global G3BP1 abundance prior to stress: G3BP1 levels decrease during HeLa cell senescence as measured by both immunofluorescence (Supp Fig. 3A) and western blot (Supp Fig. 3B), whereas fibroblasts show only a modest ∼3% increase (Supp Fig. 3N). Thus, neither reduced nor mildly increased cellular G3BP1 levels explain the early nucleation of Alt-SGs and higher enrichment in SGs.

Age-related increase in SG localization of nuclear aggregation-prone RBPs, such as FUS and TDP43, has been reported (Mokin et al. 2024; Tang et al. 2024; Koike et al. 2021), along with enhanced cytoplasmic localization of these aggregation-prone proteins (Higelin et al. 2016; Rhine et al. 2025). In contrast, hnRNPA1, a nuclear RBP also enriched in SGs shows reduced expression with age (Ziaei et al. 2012; Wang et al. 2016; Jia et al. 2019). Expressions of different SG-RBPs are reported to be variable with aging, hence we examined the localization of nuclear RBPs within alt-SGs during senescence. Consistent with the increased G3BP1 enrichment in Alt-SGs, we observed elevated levels of FUS (Fig. 3D, E), TDP43 (Fig. 3G, H), and hnRNPA1 (Supp Fig. 3I, J) in Alt-SGs compared with SGs in control cells. A similar trend was observed for eIF3η, a cytoplasmic translation-initiation complex component (Supp Fig. 3I, L). RBP-to-RNA ratio analysis further showed that, like the increased G3BP1/RNA ratio, the FUS/RNA (Fig. 3F), TDP43/RNA (Fig. 3I), hnRNPA1/RNA (Supp Fig. 3K), and eIF3η/RNA ratios (Supp Fig. 3M) were all elevated in Alt-SGs relative to control SGs. Notably, the FUS/RNA ratio was also increased in Alt-SGs of aged human fibroblasts (Supp Fig. 3O, R). Again, the increased enrichment of RBPs in alt-SGs did not directly correlate with their overall cellular concentrations. The cellular hnRNPA1 levels decreased in senescent HeLa cells (Supp Fig. 3E, F), whereas the FUS (Supp Fig. 3C, D), TDP43 (Supp Fig. 3G, Fibroblast-Supp Fig. 3P), and eIF3η (Supp Fig. 3H) were elevated in senescent cells, although enrichment of all these RBPs increased in Alt-SGs.

SGs contain ∼300 distinct proteins (Jain et al. 2016) and as all five SG-RBPs we examined showed higher protein-to-RNA ratios in Alt-SGs than in control SGs, we predicted that Alt-SGs may have an overall higher protein concentration. To measure total protein within SGs, we labeled all accessible primary amines using fluorescent amine-reactive NHS esters (Ward et al. 2017; Smolka et al. 2005) after 1 h of oxidative stress, which induces micron-sized SGs (Fig. 3J, K). NHS labeling did not show specific enrichment in SGs—consistent with SGs having cytoplasm-like density (Schlüßler et al., 2022), but strongly labeled the nucleolus (yellow arrows, Fig. 3L), a condensate with high protein density (McCall et al. 2020), confirming assay sensitivity. Despite reported proteostasis defects in senescent cells, total cellular protein content showed a modest 12% decrease, as measured by mean NHS intensity (Fig. 3L, M), consistent with reduced translation in aged cells (Khatir et al. 2023; Kim and Pickering 2023). We next quantified NHS intensity within and outside SGs using G3BP1 and polyA RNA as markers. Although senescent cells had lower total protein content, the protein-to-RNA ratio within Alt-SGs was ∼29% higher than in control SGs (Fig. 3N), indicating a higher effective protein concentration. Because NHS esters label only one terminus of each protein and are insensitive to protein length, these values provide a qualitative measure of protein abundance. Together, these results show that aged cells maintain an elevated protein-to-RNA ratio within Alt-SGs, which may contribute to their altered material properties and impaired dissolution on stress removal.

### Cellular RNA concentration decreases in aging

SGs sequester diverse cellular RNAs and RBPs (Khong et al. 2017), and reduced total RNA leads to smaller or altered SGs (Burke et al. 2020; Decker et al. 2022). During senescence, global RNA synthesis declines (Payea et al. 2024; Gyenis et al. 2023), resulting in reduced mRNA production (Park and Buetow 1990). Because Alt-SGs were depleted of polyA-tailed mRNAs, we asked whether reduced total cellular RNA underlies this phenotype. We quantified total RNA using three complementary approaches: 1) dye-accessible RNA using the RNA-selective SYTO dye (Wu et al. 2020); 2) newly synthesized RNA using 24-hour EU metabolic labeling followed by Click chemistry; 3) total extracted RNA analyzed by gel electrophoresis. In senescent HeLa cells, dye-accessible RNA (Fig. 4A, B), EU-labeled RNA (Fig. 4G, H), and polyA-tailed mRNA measured by FISH (Fig. 4E, F) were all significantly reduced prior to SG induction. Total RNA gels also showed a global decrease across RNA size classes (Fig. 4C), with 28S rRNA reduced by ∼27% (Fig. 4D). Aged mouse fibroblasts similarly exhibited decreased EU-labeled RNA (Fig. 4I, J), although polyA-mRNA levels remained comparable to young controls (Supp. Fig. 4E). In aged human fibroblasts, both EU-labeled RNA and polyA-mRNA (Supp. Fig. 4B, C) were reduced, while dye-accessible RNA remained unchanged (Supp. Fig. 4A). Across all cellular senescence and aging models, polyA-tailed mRNA and/or total EU-labeled RNA levels were consistently decreased. To test whether this reduction occurs at the tissue level, we examined *Drosophila* brains and found lower dye-accessible RNA levels in aged neuronal cell bodies than in young brains (Fig. 4K, L).

**Figure 4:**
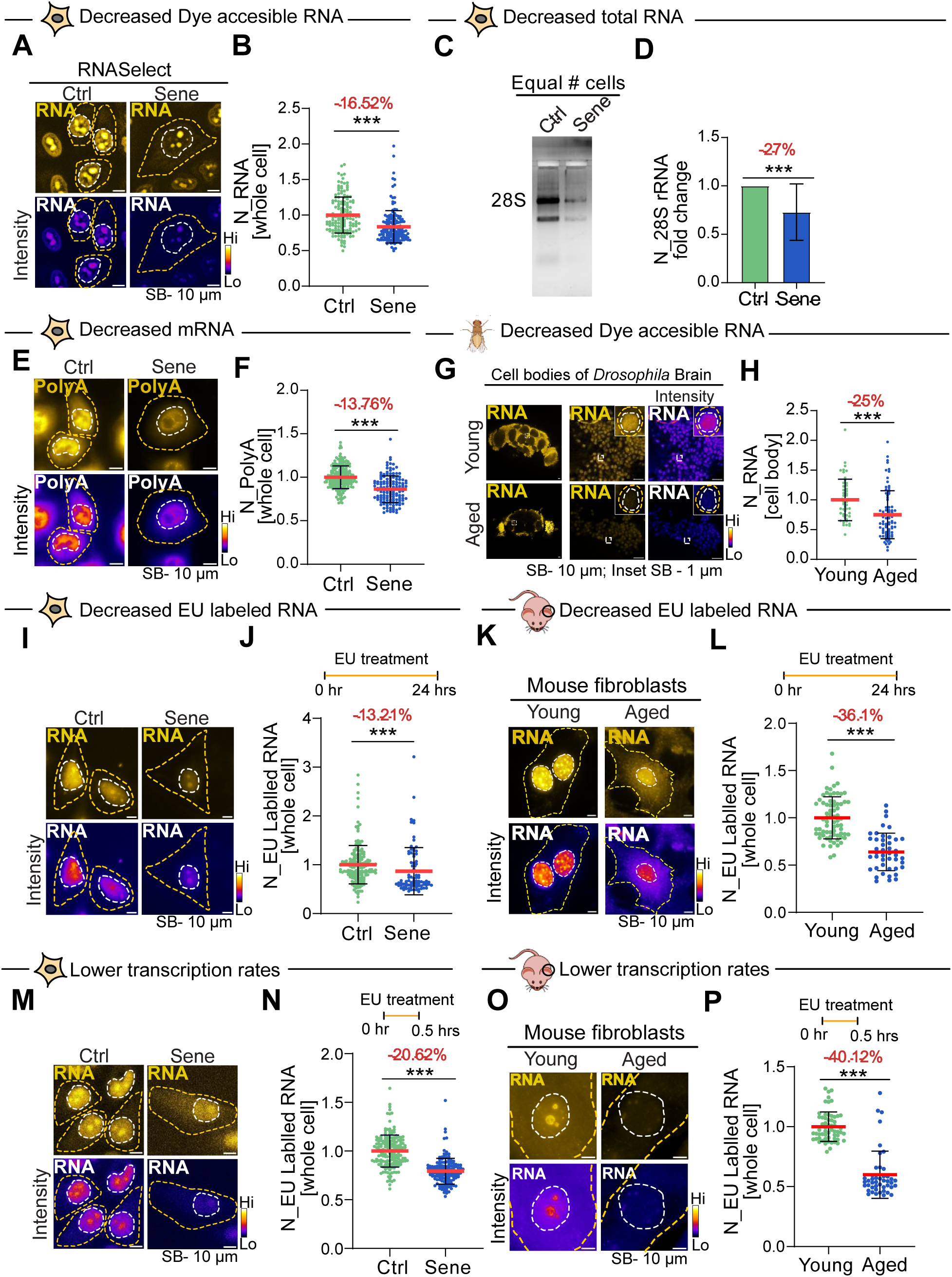
Cellular RNA concentration decreases in aging. (A) Detection of RNA using RNA-specific dye RNASelect in HeLa cells. (B) Quantification of cellular RNA content expressed as mean RNASelect intensity per cell. For each condition, ≥150 cells were analyzed. (C) Gel electrophoresis of total RNA isolated from 1 million control and senescent HeLa cells. (D) Bar graph showing 28S rRNA band intensity from RNA gels under control and senescent conditions. (E) Detection of polyA-tailed RNA under physiological conditions in HeLa cells. (F) Quantification of mean polyA levels in control and senescent cells. For each condition, ≥140 cells were analyzed. (G) Brains and neuronal cell bodies from young (3-day-old) and aged (40-day-old) *Drosophila* stained with an RNA-specific dye. White boxes mark regions enlarged in insets, highlighting nuclear and cytoplasmic RNA signals. (H) Quantification of total RNA levels. For each condition, >100 neuronal cell bodies were analyzed cumulatively from at least 18 brains. Metabolic labeling of RNA after 24 h EU exposure in (I) control and senescent HeLa cells and (K) mouse fibroblasts. Quantification of EU-labeled RNA levels in (J) control and senescent HeLa cells and (L) young and aged mouse fibroblasts. For each condition, ≥85 cells were analyzed. Transcription rates in HeLa cells and mouse fibroblasts. Metabolic labeling of nascent RNA following 30-min EU exposure in (M) HeLa cells, and in (O) mouse fibroblasts. Quantification of nascent RNA synthesis in (N) HeLa cells and (P) mouse fibroblasts. For each condition, 145 cells were analyzed. All data represent mean ± SD unless specified and are normalized to mean intensity values of respective conditions. Statistical significance determined by Mann–Whitney U test (*P < 0.05, **P < 0.01, ***P < 0.001).

To determine the cause of the age-associated reduction in cellular RNA, we quantified transcription by exposing cells to EU for 30 minutes. Senescent HeLa cells showed a ∼20.6% decrease in transcription (Fig. 4M, N), while aged mouse fibroblasts exhibited an even stronger ∼40.1% reduction (Fig. 4O, P). In contrast, aged human fibroblasts exhibited reduced total RNA (Supp. Fig. 4B, C) without a significant decline in transcription (Supp. Fig. 4D), suggesting the presence of additional regulatory mechanisms. These observations indicate that both reduced transcription and altered RNA stability contribute to global RNA loss in different aging models.

We next measured RNA degradation by tracking 30-minute EU-pulse-labeled RNA following Actinomycin D (Act D) treatment, which blocks new transcription. In senescent HeLa cells, RNA degradation rates were comparable to controls (Supp. Fig. 4F–H), indicating that transcriptional reduction is the main cause of RNA decline in these cells. Overall, across multiple senescence and aging systems, transcription and RNA degradation rates vary, but they all converge on a consistent reduction in global cellular RNA levels. Importantly, this widespread decrease in RNA shifts the RNA–RBP balance within SGs, contributing to the formation of Alt-SGs. Thus, total cellular RNA loss in aging fundamentally alters SG composition and drives the emergence of the Alt-SG phenotype.

### RNA drives the liquid-like properties of RNA-RBP condensates

Our results show that cellular aging alters both the RNA and RBP composition of SGs. Elevated RNA concentration (Maharana et al. 2018; Banerjee et al. 2017) and greater RNA single-strandedness (Maharana et al. 2022) further promote condensate dissolution. RBP alterations can also reshape SG RNA content (Mariani et al. 2024). Given the interdependence of RNA and RBPs, it is challenging to attribute SG alterations in senescent cells to either component alone. Because SG behaviour under variable RNA abundance has not been systematically examined, we used minimal and complex in vitro reconstituted SG systems to dissect how RNA concentration influences RBP stoichiometry and dynamics.

We purified G3BP1-GFP from bacteria (Supp. Fig. 5A, B), which forms condensates with or without RNA (Yang et al. 2020; Guillén-Boixet et al. 2020). Simple G3BP1 or homogenous condensates reconstituted with HeLa total RNA (Fig. 5A) showed no change in mean G3BP1 concentration within condensates (Fig. 5B, C) or in the fraction of G3BP1 partitioned (Fig. 5D). As alt-SGs in aged cells are RNA-depleted but enriched for G3BP1 and other SG-RBPs, we next examined SG-like or heterogeneous condensates formed by adding purified G3BP1-GFP to cell lysates (Fig. 5E), which recapitulate native RNA and RBP diversity (Wheeler et al. 2016; Freibaum et al. 2021). Introducing additional total RNA markedly reduced G3BP1 enrichment (Fig. 5F, G) and phase separation of G3BP1 in these SG-like condensates (Fig. 5H), demonstrating that complex condensates respond differently to RNA than single-component G3BP1 condensates.

**Figure 5:**
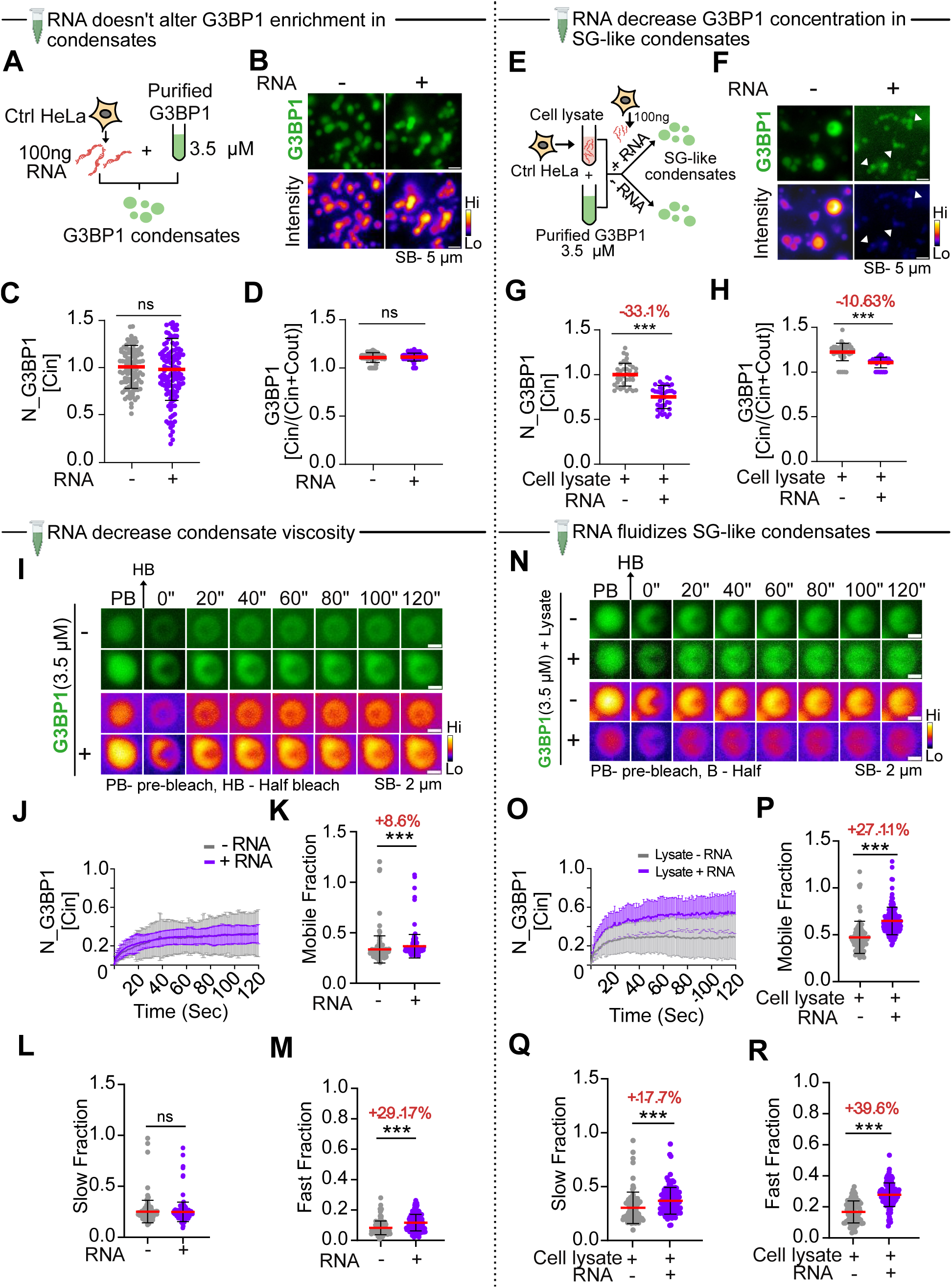
RNA drives the liquid-like properties of RNA-RBP condensates. (A) Schematic of in-vitro reconstitution of GFP-tagged G3BP1 condensates formed with purified G3BP1 in the presence or absence of total RNA from HeLa control cells. (B) Representative G3BP1 condensates formed with and without RNA. (C) Quantification of G3BP1 condensates, shown as mean intensity and (D) fraction of condensed protein. Cin denotes the fraction of total protein in the condensate and Cout denotes the fraction of total protein in the soluble phase outside the condensate. (E) Schematic of in-vitro reconstitution of SG-like condensates induced by adding purified GFP-tagged G3BP1 to HeLa cell lysate, with or without additional total RNA. (F) Representative SG-like condensates marked by G3BP1-GFP with and without RNA. (G) Quantification of SG-like condensates, shown as mean G3BP1 intensity and (H) fraction of condensed protein. (I) Montage of Half-FRAP of G3BP1 condensates prior to bleaching (PB) and after half-bleach (HB). Selected granules were half-bleached for 50 milliseconds, and fluorescence recovery was recorded for 120 seconds. (J) Mean FRAP recovery curves for half-bleached G3BP1-condensates; error bars represent standard deviation for at least *n = 200* curves per condition. (K) Mobile fraction of G3BP1-condensates calculated from individual recovery curves. (L) Quantification of slow fraction (diffusion from solution) and (M) fast fraction (intracondensate rearrangement) to overall mobile fraction as shown in K. (N) Montage of Half-FRAP of SG-like condensates prior to bleaching (PB) and after half-bleach (HB). Selected granules were half-bleached for 50 milliseconds, and fluorescence recovery was recorded for 120 seconds. (O) Mean FRAP recovery curves for half-bleached SG-like-condensates; error bars represent standard deviation for at least *n = 50* curves per condition. (P) Mobile fraction of SG-like condensates calculated from individual recovery curves. (Q) Quantification of slow fraction (diffusion from solution) and (R) fast fraction (intracondensate rearrangement) to overall mobile fraction as shown in O. All data represent mean ± SD unless specified and are normalized to mean intensity values of respective conditions. Statistical significance determined by Mann–Whitney U test (*P < 0.05, **P < 0.01, ***P < 0.001).

To assess the material properties of condensates, we performed half-FRAP (Fig. 1K). Simple G3BP1 condensates showed modest changes: raw recovery curves appeared similar (Fig. 5I, J), but fitted half-FRAP curves (Supp Fig. 5C) and quantitative analysis revealed an increased mobile fraction (Fig. 5K) driven solely by the fast component (Fig. 5L, M), with a longer fast-fraction timescale upon RNA addition (Supp. Fig. 5D, E), indicating mild fluidization. In contrast, SG-like condensates exhibited pronounced RNA-dependent changes (montage- Fig. 5N; raw curves- Fig. 5O; fitted half FRAP curves- Supp Fig. 5F) as quantified by the mobile fraction (Fig. 5P), both fast and slow fractions increased (Fig. 5Q and R), with a reduced slow-fraction timescale (faster exchange from solution) and an increased fast-fraction timescale (Supp. Fig. 5F–H). Thus, RNA has a modest effect on purified homogenous G3BP1 condensates but induces substantial fluidization in complex heterogenous SG-like condensates. Overall, our reconstitution experiments demonstrate that RNA concentration is a key determinant of SG fluidity.

### Increase cellular RNA reverses persistent SGs

Our in vitro and ex-cellulo reconstitution assays of SG-like condensates showed that reduced RNA concentration alters SG properties and likely drives the Alt-SG phenotype in aging. Because SGs sequester diverse RNAs (Khong et al. 2017; Van Treeck et al. 2018; Glauninger, Bard, Hickernell, et al. 2025), and total cellular RNA decreases in parallel with Alt-SG RNA, we predicted that increasing cellular RNA would similarly increase Alt-SG RNA. Aged cells typically exhibit reduced nucleotide pools (Aird et al. 2013; Nikiforov and Shewach 2017), and exogenous nucleosides can restore these pools (Aird et al. 2013; Delfarah et al. 2019). We induced senescence with 2 days of BrdU treatment, supplemented the media with 30 µM of each nucleoside for the next 2 days (Fig. 6A), and measured RNA on day 4.

**Figure 6:**
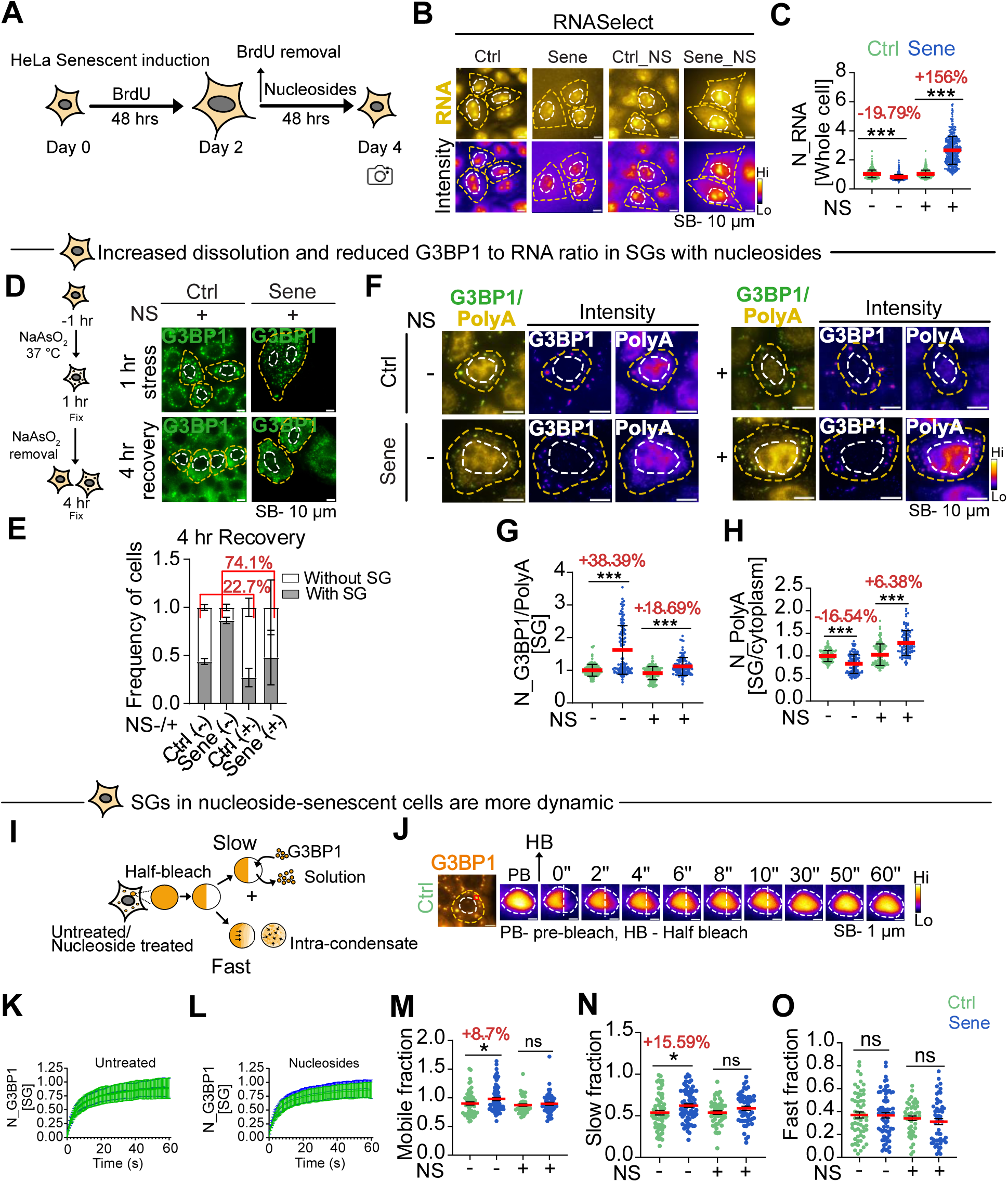
Increase cellular RNA reverses persistent SGs. (A) Schematic of senescence induction followed by BrdU removal and nucleoside supplementation (Day 2) for 48 hours (Day 4). (B) Control and senescent HeLa cells (NS⁺/⁻) stained with the RNA-specific dye RNASelect. (C) Quantification of cellular RNA content expressed as mean RNASelect intensity per cell. For each condition, ≥ 200 cells were analyzed. (D) SG dissolution in control and senescent HeLa cells (NS⁺/⁻) after 1 h of NaAsO₂ stress followed by 4 h recovery, detected with antibodies against endogenous G3BP1. (E) Bar graph shows the normalized frequency of cells with or without SGs at 4 hr stress recovery. (F) Control and senescent HeLa cells (NS⁺/⁻) stained for G3BP1 and probed for polyA-tailed mRNA using fluorescent oligo(dT) FISH after 1 h of 0.25 mM NaAsO₂ stress. (G) Ratio of G3BP1 to polyA within SGs, calculated as mean G3BP1 intensity divided by mean polyA intensity per cell. For each condition, atleast 70 cells were analyzed. (H) Quantification of polyA enrichment in SGs, expressed as the ratio of mean polyA intensity in SGs to cytoplasm. (I) Half-FRAP of SGs tagged with G3BP1-mCherry (NS⁺/⁻), formed after 1 h of NaAsO₂ treatment at 37 °C. The schematic illustrates two mechanisms of fluorescence recovery: (i) slow diffusion of SG components from the cytosol into the condensate and (ii) rapid molecular rearrangement within the condensate. (J) Montage of half-frapped G3BP1-mCherry–labeled SG induced by 1 hour of NaAsO₂ stress used for calculation of mean recovery curve. Images show SGs before bleaching (PB) and after a half-bleach (HB). Selected granules were half-bleached for 50 milliseconds, and fluorescence recovery was recorded for 60 seconds. Mean FRAP recovery curves (K) untreated and (L) nucleoside-treated plotted by quantifying normalized mean G3BP1 intensity in half-bleached SGs (mean ± SD; ≥60 curves per condition). (M) Mobile fraction of SGs derived from individual recovery curves. Quantification of (O) slow fraction and (P) fast fraction (intracondensate rearrangement) contributing to overall mobility. All data represent mean ± SD unless specified and are normalized to mean intensity values of respective conditions. Statistical significance determined by Mann–Whitney U test (*P < 0.05, **P < 0.01, ***P < 0.001).

Nucleoside supplementation (NS) did not change dye-accessible RNA in young cells (Fig. 6B, C), likely because their sufficient endogenous nucleotide pools suppress nucleoside import (Boswell-Casteel and Hays 2017; Pastor-Anglada and Pérez-Torras 2018). In senescent cells, NS fully rescued the RNA deficit and further increased total RNA compared to untreated cells (Fig. 6B, C). NS also increased polyA-tailed mRNA, which had declined in senescence (Fig. 4E, F; Supp Fig. 6A), and elevated total RNA in primary aged mouse fibroblasts (Supp Fig. 6B). Because reduced transcription contributed to RNA loss in aged cells, we measured nascent RNA synthesis after a 30-min EU pulse (Supp Fig. 6C). NS increased transcription rates and reduced the difference in EU intensities between control and senescent cells. Senescent cells degraded ∼22% of RNA in 2 h compared to ∼12% in controls; NS reduced degradation in senescent cells to ∼2% while leaving control cells unchanged (Supp Fig. 6D). Thus, NS increased cellular RNA by boosting transcription and reducing degradation.

We next asked whether NS improved SG and Alt-SG dissolution after stress removal. After 4 h of recovery, 56% of control cells lacked SGs, whereas only 13% of senescent cells did. NS increased recovery to 72% in controls and 52% in senescent cells—an ∼74% improvement (Fig. 6D, E). Because high RBP/RNA ratios coincided with persistent Alt-SGs (Fig. 3) and in vitro data showed that low RBP/RNA enhances condensate dynamics, we examined whether NS altered this ratio. Alt-SGs showed ∼38% higher G3BP1/polyA ratios than SGs (Fig. 6F, G). NS reduced this difference to 18%, making Alt SGs more similar to control SGs. A ∼6% increase in polyA RNA within Alt-SGs primarily drove this shift (Fig. 6H). These results demonstrate that increasing cellular RNA elevates condensate RNA content and enhances SG dissolution during recovery.

As persistent Alt-SGs exhibit higher viscosity (Fig. 1L–O), we examined how NS alters SG material properties. Before NS, raw and fitted half-FRAP curves showed similar recoveries in control SGs and Alt-SGs (Fig. 6K, Fig. 6I, J). After NS, senescent SGs displayed slightly improved recovery relative to controls (Fig. 6L; Supp Fig. 6E, F). NS equalized the mobile fractions between SGs and Alt-SGs (Fig. 6M). Without NS, Alt-SGs recovered mainly through slow, diffusion-limited exchange, evident from a ∼16% increase in slow fractions (Fig. 6N). NS made both the slow (Fig. 6N) and fast (intradroplet mixing) fractions (Fig. 6O), as well as their respective timescales (Supp Fig. 6G, H), which were similar between SGs and alt-SGs. Thus, increasing global RNA levels in senescent cells rejuvenates Alt-SG material properties, reduces their viscosity, and enhances SG dissolution during stress recovery.

### Restored RNA metabolism rejuvenates senescent phenotypes by reversing protective SGs

SGs represent one of several RNA–RBP condensates, raising the possibility that NS treatment—which elevates global RNA concentration—may broadly influence condensate behavior and the senescence phenotype. To test this, we quantified senescence-associated markers following NS treatment. NS exposure reduced senescence-associated β-galactosidase activity (Fig. 7A, B) and lowered the expression of canonical senescence markers (Fig. 7C). Live-cell imaging of SG dissolution after stress removal revealed that ∼37% of senescent cells underwent cell death by bursting during the 3-hour recovery period from oxidative stress (Fig. 7D, E), whereas no control cells died under the same conditions. We therefore examined whether NS treatment modulates this stress fluctuation-induced death (Fig. 7I). Indeed, NS treatment reduced the proportion of dying senescent cells from 37% to 20% during recovery (Fig. 7F, J). Notably, cells that succumbed to death consistently exhibited smaller total SG areas, regardless of NS treatment (Fig. 7G, K), while G3BP1 enrichment within SGs remained unchanged (Fig. 7H, L).

**Figure 7:**
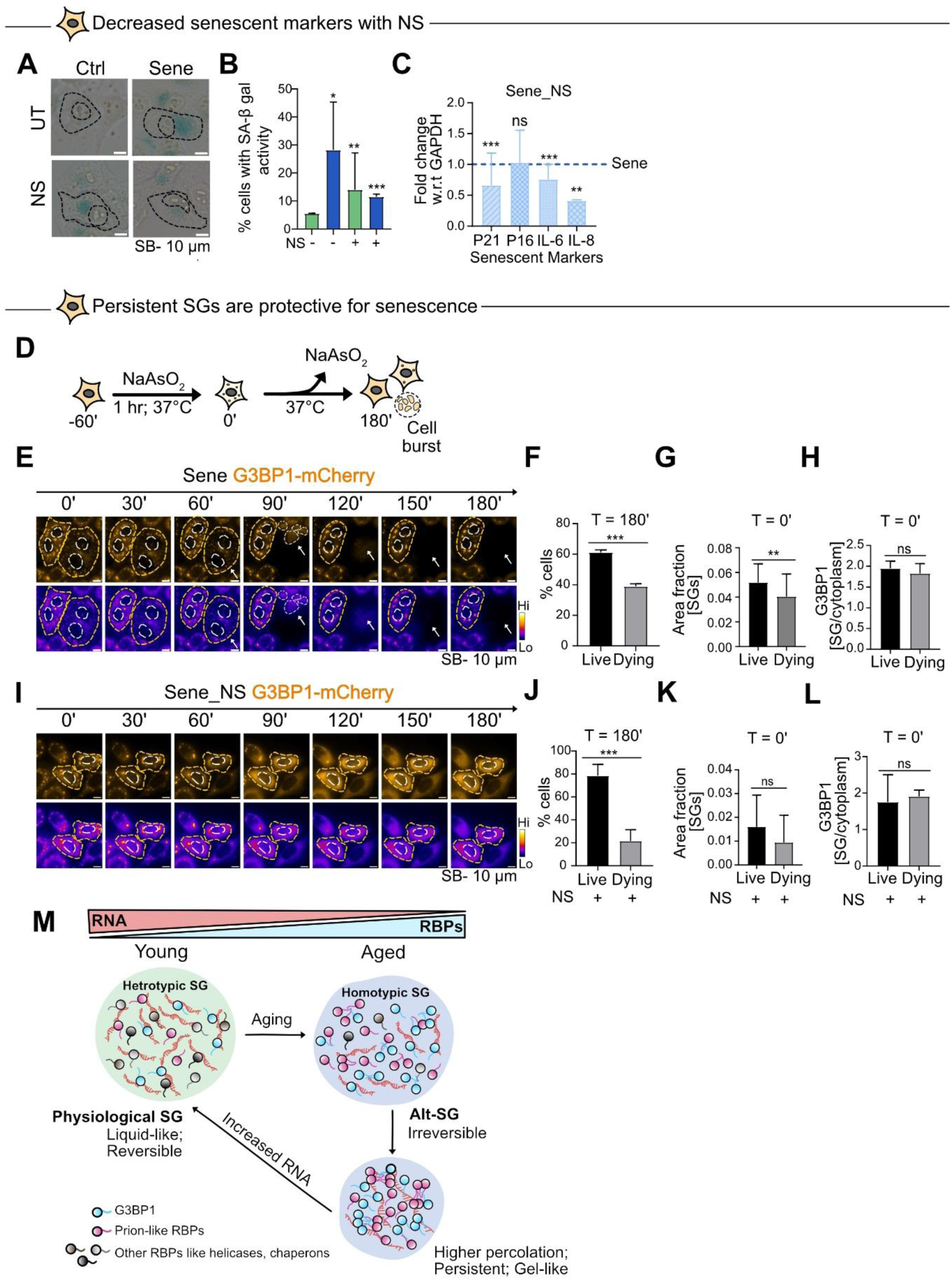
Restored RNA metabolism rejuvenates senescent phenotypes by reversing protective SGs. (A) Montage of control and senescent (NS⁺/⁻) cells showing SA-β-gal staining. (B) Quantification includes the percentage of SA-β-gal-positive cells, (C) mRNA expression levels of P21, P16, IL-6, and IL-8 in NS-treated senescent versus senescent conditions. (D) HeLa cells expressing G3BP1-mCherry were stressed with NaAsO₂ (60 min, 37 °C) to induce SGs; following stress removal, SG dissolution was monitored by live-cell imaging for 180 min, where cell bursts were captured. (E) Montage of SGs in G3BP1-mCherry expressing senescent HeLa cells following NaAsO₂ removal. 0’ shows SGs formed after 1 h of stress, prior to stress withdrawal. The white arrows in the subsequent panels depict cell bursts during the indicated time points. Quantification of (F) percentage of live and dying cells at 180 min, (G) SG area fraction in live and dying senescent cells, and (H) mean G3BP1 enrichment within SGs per live and dying senescent cells after stress, 0’. For each condition, atleast 20 cells were analyzed across atleast 15 independent time-lapse movies. (I) Montage of SGs in G3BP1-mCherry expressing NS-treated senescent HeLa cells following NaAsO₂ removal. 0’ shows SGs formed after 1 h of stress, prior to stress withdrawal, subsequent panels depict SG dissolution during the indicated time points. Quantification of (J) percentage of live and dying cells at 180 min, (K) SG area fraction in live and dying NS-treated senescent cells, and (L) mean G3BP1 enrichment within SGs per live and dying NS-treated senescent cell after stress, 0’. For each condition, atleast 50 cells were analyzed across atleast 10 independent time-lapse movies. All data represent mean ± SD unless specified and are normalized to mean intensity values of respective conditions. Statistical significance determined by Mann–Whitney U test (*P < 0.05, **P < 0.01, ***P < 0.001).

Together, these findings demonstrate that NS treatment attenuates the senescent phenotype and mitigates stress fluctuation-induced cell death. Moreover, the persistence of SGs appears to confer a protective advantage, enhancing the adaptability and survival of senescent cells under fluctuating stress conditions.

## Discussion

Aging involves pervasive remodelling of cellular organization beyond classical organelle dysfunction. While membrane-bound organelles have been extensively implicated in age-related decline, far less is known about how aging reshapes the formation, composition, and material properties of membraneless RNA-RBP biomolecular condensates-the reaction hubs that coordinate gene expression and stress responses. Here, we show that cytoplasmic SGs, which comprise translationally relevant RBPs and aggregation-prone RBPs in addition to RNAs, in aged cells adopt altered material states-**alt-SGs**-that are persistent and slow to dissolve after stress removal. Mechanistically, alt-SGs arise due to an **increase in viscosity** due to elevated **RBP/RNA ratios** within condensates, a direct consequence of diminished RNA metabolism and reduced global RNA concentrations in aged cells. By systematically decoupling the effects of RNA and RBPs across cells, lysates, and minimal reconstitutions, we demonstrate that condensate RNA concentration is sufficient to reprogram RBP composition and phase behaviour, thereby dictating condensate dynamics and reversibility.

SGs form upon translation inhibition by sequestering ribosome-released mRNAs and numerous translation-associated RBPs. Their physiological function depends on rapid molecular exchange with the cytoplasm so that SGs remain sensitive to translation levels. Our results show that SGs in senescent cells are persistent and hence could be insensitive to the translation requirements of the cells. Further we found that **heterogeneous**, SG-like condensates nucleated by G3BP1 were substantially **more dynamic** than **minimal**, 1-2 component G3BP1 condensates (Fig. 5K, P). Adding RNA only marginally fluidized homogeneous G3BP1 droplets but markedly increased the dynamics of SG-like heterogeneous condensates. SGs of young cells-which dissolved readily upon stress removal-were more liquid-like and displayed lower RBP/RNA ratios than persistent alt-SGs, which were enriched in RBPs relative to RNA (Fig. 3A-I).

Our data support two non-mutually exclusive mechanisms by which **condensate RNA concentrations** promote SG fluidity and reversibility: **(i) RNA-driven heterogeneity decreases percolation.** SGs are highly heterogeneous assemblies where G3BP1/2 (Jain et al. 2016; Markmiller et al. 2018; Marmor-Kollet et al. 2020) represent only a small fraction of ∼300 RBPs (Jain et al. 2016) coexisting with hundreds of RNA species (Khong et al. 2017; Demeshkina and Ferré-D’Amaré 2025). Multivalent G3BPs-other SG RBP and RBP-RNA interactions (Guillén-Boixet et al. 2020; Van Treeck et al. 2018) diversify the network of interactions within SGs. RNA lowers the critical concentration for phase separation of many RBPs (Maharana et al. 2018; Joseph et al. 2021), thereby **increasing compositional heterogeneity** (Fig 7M). When condensates are relatively homogeneous or enriched in aggregation-prone RBPs (e.g., FUS, TDP-43, hnRNPA1), short-range high-affinity interactions promote gelation/solidification (Emmanouilidis et al. 2024; Feng et al. 2026; Guo et al. 2026). Increased heterogeneity reduces the effective concentration of aggregation-prone RBPs in SGs and **limits percolation**, maintaining fluidity (Alshareedah et al. 2019). **(ii) RNA-dependent recruitment of helicases and chaperones.** SG RNAs recruit RNA helicases (e.g., DDX3X, DHX36) (Samir et al. 2019; Cheng et al. 2025) and RNA-interacting chaperones (Hsp70, HSPB8, Hsp90, HspBP1, eIF4A2) that **reduce persistence and increase exchange** (Ganassi et al. 2016; Mediani et al. 2021; Mahboubi et al. 2020; Jongjitwimol et al. 2016). Thus, higher condensate RNA concentrations simultaneously **diversify the interaction network** and **enrich remodelling factors**, jointly enhancing condensate dynamics (Fig 7M).

We find that condensate RNA concentrations are constrained by the absolute cellular RNA pool. Across aging models in human, mouse, and fly tissues, total RNA abundance declines (Fig. 4), primarily due to reduced transcriptional output (Fig. 4M–P) (Gyenis et al. 2023; Tahoe et al. 2004). Aged cells also experience nucleotide insufficiency, further limiting transcription. Supplementing nucleosides restored nascent transcription and elevated global RNA levels in senescent cells, but not in young cells (Fig. 6B, C), likely because young cells maintain sufficient intracellular nucleoside pools and tightly regulate uptake (Boswell-Casteel and Hays 2017; Pastor-Anglada and Pérez-Torras 2018). Elevating global RNA selectively increased SG RNA content and reversed senescence-associated phenotypes (Fig. 6H, 7A–C), while leaving RNA levels unchanged in young cells (Fig. 6B, C), indicating targeted normalization of age-disrupted RNA metabolism. Similar rejuvenating effects have been reported with supplementation of nucleotide-related metabolites such as NADH (H. Wang et al. 2022; Yuan et al. 2020) and glutamine (Zhou et al. 2022; Zhang et al. 2024), which act both as signaling intermediates and metabolic substrates for RNA/DNA synthesis. Together, these findings demonstrate that restoring nucleotide availability can restart transcription, elevate global RNA, and rebalance RBP/RNA ratios within condensates.

By quantifying both SG nucleation and dissolution in **live aged cells**, we identify kinetic hallmarks of alt-SGs. Aged cells nucleate SGs more efficiently (Fig. 1), yet total SG quantity per cell can be lower (Fig. 1D) (Dubinski et al. 2023). Critically, dissolution is slower, yielding **more persistent SGs** (Fig. 1F-J) in aging. Because organisms experience recurrent physiological stress, impaired dissolution may underlie the **spontaneous SGs** we observe in the aged mouse and fly brains (Fig. 2J-R). Aged SGs were often **smaller** (Supp. Fig. 2E-G), consistent with rapid nucleation and **reduced fusion** (Supp. Fig. 1D). Extensive FRAP measurements revealed **increased intracondensate viscosity** with age (Fig. 1K-P), implicating physical-state changes in persistence of SGs. Notably, an **increased RBP:RNA ratio** within SGs was the most consistent signature across different aging models and model organisms. In aged cells and in SG-like in vitro condensates, **decreased RNA content** increased the absolute **enrichment of G3BP1 and hnRNPA1 within SGs** (Fig. 2A, B; Supp Fig. 3I, J), despite their global decline with age (Supp Fig. 3A, E), highlighting RNA as a **primary determinant** of RBP partitioning. Since aging and neurodegeneration are associated with elevated cytoplasmic FUS and TDP-43 (Higelin et al. 2016; Tang et al. 2024; Rhine et al. 2025), we propose that low-RNA SGs preferentially enrich for **aggregation/phase-prone RBPs**, amplifying high-affinity interactions and persistence. Total-protein labeling further confirmed that aged SGs contain more protein per unit of **RNA** (Fig. 3J-N), reinforcing a shift toward protein-dense networks.

More broadly, condensate formation is a **biophysical process** governed by multivalent, weak interactions and, for many RNA-RBP condensates, is **insensitive to RNA sequence** but highly sensitive to **total RNA concentration** (Maharana et al. 2018; Banerjee et al. 2017). Yet aging- and disease-associated defects in RNA metabolism are typically quantified as **relative** expression changes, which obscure **absolute** RNA amounts per cell. Our findings argue that reduced **global RNA concentration** is sufficient to alter the composition and material state of multiple RNA-RBP condensates beyond SGs, including **nuclear speckles** and the **nucleolus** (unpublished observations), with implications for splicing, ribosome biogenesis, and proteostasis. Further, protein-dense networks in the condensates also increase the probability of forming more structured assemblies of aggregation-prone proteins like TDP43 and FUS within the aging SGs (Fig. 7M, lower right) (Yan et al. 2025).

We propose that aging-driven **RNA scarcity** elevates condensate **RBP/RNA ratios**, reducing heterogeneity, weakening remodelling capacity, and **shifting SGs toward persistent, protein-dense states**. Restoring RNA biosynthesis in aging by **metabolic supplementation** reconstitutes condensate RNA levels, **rebalances composition**, and **rejuvenates condensate dynamics**. RNA metabolism—and specifically the absolute concentration of RNA-acts as a central rheostat that sets the composition, material state, and reversibility of RNA-RBP condensates in aging cells.

## Lead Contact and Materials Availability

Further information and requests for resources and reagents should be directed to and will be fulfilled by the lead contact, Shovamayee Maharana.

This study did not generate new, unique reagents.

## Acknowledgements

We thank the members of Dr Shovamayee Maharana’s and Dr Anand Srivastava’s laboratory for insightful discussions. We are grateful to the IISC imaging facility, the departmental Tecan plate reader facility, the spectrophotometer facility, the PCR facility, the NCBS fly facility, and the mouse facilities at both IISC and IIT Kanpur for their support. We acknowledge Brishti Kundu, a dissertation student, for assistance with live-cell SG dissolution imaging experiments. We also thank Dr Baskar Bakthavatchalu, Dr Anand Srivastava, and Dr Sangeetha Balasubramanian for valuable discussions.

S.M. was supported by the DBT–WT India Alliance Intermediate Fellowship (Grant No. IA/I/21/2/505944), the CSIR Aspire (Grant No. 37WS(0079)/2023-24/EMR-II/ASPIRE), and the Infosys Young Investigator Award. A.R. was supported by the DBT Junior Research Fellowship Programme; S.S.R. by the UGC Junior and Senior Research Fellowship Programmes; A.S. by the MOE (Ministry of Education) Fellowship Programme; and P.G.D. by the CSIR Junior and Senior Research Fellowship Programmes. T.P. was supported by DBT–WT India Alliance grant awarded to Dr Shovamayee Maharana. N.B. was supported by the Longevity India Initiative, IISC program. S.G. received support from the Indian Council of Medical Research (Grant No. 5/3/8/20/2019-ITR), Government of India; the Science and Engineering Research Board (SERB) (Grant No. JCB/2022/000007), Government of India; and jointly with R.P. from SERB (Grant No. CRG/2020/001371). A.C. was supported by DBT–WT India Alliance (Grant No. IA/I/19/1/504286) grant awarded to Dr Baskar Bakthavatchalu.

## Author contributions

S.M. conceived the project, designed the experiments, and supervised the study. A.R. conceived the project, designed and conducted the experiments, and performed data analysis. S.G. contributed to the conceptual development of the mouse experiments. T.P. performed and analyzed the in vitro experiments, while S.S.R. carried out and analyzed the half-FRAP experiments. N.B. conducted experiments on *Drosophila* Malpighian tubules and analyzed SG fusion in cell lines. R.P. performed fibroblast isolation and brain immunohistochemistry in mouse experiments, and A.S. performed and analyzed Western blot experiments. A.C. carried out immunohistochemistry in *Drosophila* brain experiments. P.G.D. developed custom MATLAB (R2023a) codes. All authors contributed to data analysis and interpretation, and the manuscript was written by A.R. and S.M.

## Methodology

### Cell culture

HeLa Kyoto cells (a kind gift from Dr Sachin Kotak, Indian Institute of Science, Bengaluru, India) and HeLa G3BP1-mCherry BAC stable cells (a kind gift from Prof. Anthony Hyman, MPI-CBG, Dresden; Poser et al. 2008) were maintained in DMEM supplemented with 5% fetal bovine serum (FBS) and 1% penicillin–streptomycin at 37 °C in a humidified incubator with 5% CO₂. Cells were grown in glass flasks (Corning™ PYREX™ Milk Dilution Bottles with Screw Cap) and passaged by trypsinization once they reached ∼80% confluency. For BAC cell lines, the medium was additionally supplemented with blasticidin (5 µg/ml).

Primary human skin fibroblasts (derived from young (GM01652) and aged donors (GM08401), Coriell Institute) were cultured in Eagle’s Minimum Essential Medium (EMEM) supplemented with 15% FBS and 1% penicillin–streptomycin under identical incubation conditions (37 °C, 5% CO₂).

### Mouse fibroblast isolation and culture

Primary fibroblasts were isolated from ear punches of two-month-old and eighteen-month-old C57BL/6 mice using a modified version of the protocol described by Upadhyay et al. 2017. Briefly, 1–2 mm ear punches were collected, mechanically minced in 1× PBS, and incubated with collagenase I at 37 °C in a humidified atmosphere containing 5% CO₂ for 25 minutes. The resulting cell suspension was centrifuged at 6500 rpm for 5 minutes. The pellet was then treated with 0.1% trypsin–EDTA for 15 minutes under the same incubation conditions. Following enzymatic digestion, cells were centrifuged at 1000 rpm for 5 minutes, resuspended in DMEM supplemented with 15% FBS and 1% antibiotic–antimycotic solution, and passed through a 100 µm nylon strainer. Cells were seeded onto 0.1% gelatin-coated dishes in DMEM containing 15% FBS and 1% antibiotic–antimycotic solution. Surplus cells were cryopreserved at –80 °C in freezing medium (10% DMSO, 90% FBS). Thawed cells were used within 3–4 passages after initial seeding or thawing to minimize additional cellular senescence.

### Nucleoside treatment

Previous studies have demonstrated that exogenous nucleoside supplementation can suppress the senescence phenotype and cell growth arrest (Aird et al. 2013). Based on this, we supplemented cells with nucleosides to examine RNA metabolism recovery in aged cells. Control and senescent HeLa Kyoto cells (∼50,000 cells), as well as young and aged mouse fibroblasts, were cultured on 12 mm-diameter No. 1 glass coverslips in 24-well plates. For live SG dissolution experiments, senescent HeLa G3BP1-mCherry cells were cultured on 22 mm diameter No. 1 glass coverslips, mounted in autoclaved stainless-steel live-cell imaging dishes. These cells were then treated with nucleosides (1X; ∼0.73 µg/µL cytidine, 0.24 µg/µL thymidine, 0.8 µg/µL adenosine, 0.85 µg/µL guanosine, and 0.73 µg/µL uridine) in DMEM for 48 hours at 37 °C in 5% CO₂. Following incubation, cells were subjected to stress conditions to assess SG dynamics or to quantify total RNA.

### Senescence-associated β-galacosidase assay

Senescent cells exhibit increased activity of senescence-associated β-galactosidase (SA-β-gal), which is detectable at pH 6.0 and distinct from the lysosomal β-galactosidase activity observed at pH 4.0. To assess this, we adapted the β-galactosidase assay described by Dimri et al. 1995. The assay employs the chromogenic substrate 5-bromo-4-chloro-3-indoyl β-D-galactopyranoside (X-gal), which is cleaved by β-galactosidase to yield an insoluble blue precipitate (5,5′-dibromo-4,4′-dichloro indigo). Enhanced SA-β-gal activity in senescent cells results in increased formation of this blue precipitate.

Approximately 0.2 million BrdU-treated and control cells were seeded per well in 6-well plates. After 24 h, cells were washed with PBS and fixed at room temperature for 15 minutes in 0.2% glutaraldehyde prepared in PBS. Fixed cells were then incubated overnight at 37 °C (without CO₂) in freshly prepared staining solution containing: 1 mg/ml X-gal, 40 mM citric acid/sodium phosphate buffer (pH 6.0), 5 mM potassium ferrocyanide, 5 mM potassium ferricyanide, 150 mM NaCl and 2 mM MgCl_2_. The following day, cells were washed with PBS to remove excess stain and imaged for blue precipitate formation using an inverted IX81 microscope equipped with a DP72 color CCD camera (Olympus, Japan).

### Oxidative stress induction and recovery from stress

To induce SG formation, approximately 0.05 million cells were seeded onto 12 mm round coverslips (No. 1 thickness) placed in 24-well plates. Coverslips were either mounted in stainless steel live-cell imaging chambers from GreenFocus Research Technologies (0.2 million cells in 22 mm round coverslips) or left in the 24-well plates for subsequent fixed-cell imaging. Cells were treated with 0.25 mM sodium arsenite (NaAsO₂; stock prepared in Milli-Q water) diluted in complete culture medium for 30 minutes to 2 h at 37 °C in a humidified incubator with 5% CO₂.

For primary mouse fibroblasts, young and aged cells were exposed to 1 mM NaAsO₂ in DMEM supplemented with 15% FBS and 1% antibiotic–antimycotic solution for 1 h under the same incubation conditions. Longer exposure (2 h) resulted in significant cell death and was therefore avoided.

For SG dissolution assays, control and senescent cells were first subjected to oxidative stress (1 h, 37 °C, 5% CO₂) as described above. Following treatment, the NaAsO₂-containing medium was replaced with fresh culture medium. Cells were then fixed at 1.5, 3, and 4 h after media replacement to monitor SG disassembly over time.

### Live cell imaging of stress granule nucleation

SG formation was monitored in live control and senescent HeLa G3BP1-mCherry BAC stable cells. Approximately 0.2 million cells were seeded onto 22 mm diameter No. 1 circular coverslips mounted in autoclaved stainless-steel live-cell imaging dishes and cultured overnight at 37 °C in a humidified incubator with 5% CO₂. The following day, dishes were transferred to a microscope equipped with an onstage incubator. SGs were induced by treating cells with 0.25 mM NaAsO₂ in DMEM supplemented with 5% FBS, 1% penicillin–streptomycin, and 10 mM HEPES for 2 h at 37 °C and room temperature (25 °C) without additional CO₂.

During treatment, SG formation and fusion were imaged at 60× magnification using a Nikon ECLIPSE Ti2 microscope equipped with an ORCA®-Fusion BT camera and a CoolLED PE-800 LED illumination. Images were acquired every minute for 2 h using 5% excitation intensity of a 561 nm LED and an exposure time of ∼200 ms. To capture SGs across different focal planes, a Z-stack of 9 frames with a step size of 0.4 µm was collected. Quantification of SG area fraction and G3BP1 fluorescence intensity within granules was performed using a custom-written MATLAB (R2023a) script.

### Analysis of live cell images for SG nucleation

Time-lapse Z-stack images were acquired every minute for 2 h and subsequently maximum-projected across all time points. To correct for sample drift, the max-projected stacks were aligned in Fiji using the “Linear Stack Alignment with SIFT” registration plugin. Drift-corrected stacks were then converted into multi-TIFF files for downstream analysis in MATLAB (R2023a).

Image analysis involved segmentation of both cells and SGs. From these segmented regions, quantitative parameters were extracted, including: the percentage of cells containing SGs, the SG area fraction, and G3BP1 fluorescence intensity and enrichment within SGs.

The absolute data were plotted in R (version 4.3.1) as line graphs showing mean ± standard error to depict SG dynamics. For analysis, foci smaller than 0.0049 µm² in area (corresponding to ∼250 nm diameter) were excluded. SGs were approximated as circular structures with a radius of 125 nm (area = 0.049 µm²). Cells containing ≥ 8 SGs were classified as SG-positive.

### Quantification of SG fusion events

Fusion events of SGs were quantified from time-series movies of both control and senescent cells, using data pooled from all biological replicates. To ensure unbiased analysis, the number of cells examined was kept constant, with 25 cells selected per condition. The quantified fusion events were plotted as bar graphs using R, and statistical significance was assessed using the Mann–Whitney U test.

### Live cell imaging and analysis of SG dissolution

SG dissolution was imaged in live control and senescent HeLa G3BP1-mCherry BAC stable cells. Approximately 0.2 million cells were seeded onto 22 mm diameter No. 1 circular coverslips mounted in autoclaved stainless-steel live-cell imaging dishes and cultured overnight at 37 °C in a humidified incubator with 5% CO₂.

The following day, cells were then exposed to 0.25 mM NaAsO₂ in DMEM supplemented with 5% FBS and 1% penicillin–streptomycin for 1 h at 37 °C. For dissolution assays, NaAsO₂-containing medium was replaced with fresh medium (DMEM supplemented with 5% FBS, 1% penicillin–streptomycin, and 10 mM HEPES). SG dissolution was imaged using a Nikon ECLIPSE Ti2 microscope equipped with an onstage incubator maintained at 37 °C. Images were acquired every 5 minutes for 3 h at 60× magnification, using 5% excitation intensity of a 561 nm LED and an exposure time of ∼200 ms. A Z-stack of 9 frames with a step size of 0.4 µm was collected to capture SGs across different focal planes. Quantitative analysis of SG area fraction, G3BP1 fluorescence intensity, and percentage of SG-positive cells was performed using a custom-written MATLAB (R2023a) script. The data is further normalized to 1 hr stress (time 0 of recovery) and statistically analyzed across multiple cells over time. The normalized data were plotted using R to generate the line plot with mean and standard error that depicts the dissolution of SGs.

### *In Situ* RNA hybridisation to detect polyA-tailed mRNA and immunofluorescence

Approximately 0.05 million cells were plated onto 12 mm-diameter No. 1 round coverslips in 24-well plates. Cells were fixed with 4% formaldehyde in PBS for 10 minutes at room temperature and permeabilized with 0.25% Triton X-100 in PBS for 10 minutes.

For RNA FISH, permeabilized cells were incubated for 3 h at 37 °C in a moist chamber with 125 nM 3′-biotinylated 25-mer oligo-dT primer prepared in hybridization buffer. The hybridization buffer contained 4× SSC (0.6 M NaCl, 0.1 M sodium citrate), 10% formamide, 5% dextran sulfate, 1% BSA, and 0.5 mM EDTA. Detection of biotinylated oligo-dT primers was performed by incubating cells with Cy5-streptavidin (1:5000 dilution in 2% BSA) for 2 h (Maharana et al. 2018).

For immunofluorescence on the same samples, SSC wash buffer was replaced with PBS. After hybridization, cells were washed three times in 4× SSC buffer and blocked in 2% BSA in PBS for 1–2 h at room temperature. Cells were then incubated with primary antibodies diluted in 2% BSA for 16 h at 4 °C. Following three gentle PBS washes, cells were incubated with Alexa Fluor™-conjugated secondary antibodies (1:1000 dilution in 2% BSA/PBS) for 1 h at room temperature. Nuclei were stained with 1 µg/ml DAPI in PBS for 10 minutes at room temperature. After three PBS washes, coverslips with cells were mounted in mounting medium (10% w/v Mowiol, 25% glycerol, 2.5% DABCO in 0.1 M Tris-Cl, pH 8.5) and sealed with transparent nail polish.

### Fixed cell imaging

Imaging was performed using Nikon ECLIPSE Ti2 and Leica DMI6000b microscopes equipped with a 60×/1.518 NA oil-immersion objective (type F). Fluorescent excitation was achieved using 488 nm, 561 nm, and 647 nm LED lines from the CoolLED PE-800 for GFP, mCherry, and Far-Red, respectively. Emission was collected with a bandpass filter set of maxima- 470 nm (GFP), 550 nm (mCherry), and 635 nm (Far-Red) ±30 nm.

Images were acquired using the ORCA®-Fusion BT camera at a resolution of 2304 × 2304 effective pixels, with a pixel size of 6.5 µm × 6.5 µm. Z-stacks were captured with a step size of 0.2–0.3 µm. Laser power and gain settings were kept constant across all samples.

For representation, images were processed in Fiji. Contrast and brightness were linearly adjusted and applied equally across all conditions. Intensity-based colour mapping was done using the Fire lookup table (LUT) to colour-code pixel intensity. Final images were saved in TIFF or JPEG format for presentation.

### Fixed cell SG analysis

Quantitative analysis of SGs was performed using a custom-written MATLAB (R2023a) script. Parameters measured included SG area fraction, fluorescence intensities of G3BP1, FUS, TDP43, hnRNPA1, eIF3η, and PolyA within SGs, as well as the percentage of SG-positive cells.

For senescent or aged samples, data were normalized to their respective young controls. For nucleoside-treated samples, normalization was performed against untreated controls. Normalized values were plotted in R (version 4.3.1) as line graphs showing mean ± standard deviation.

To assess SG size distribution, the area of individual SGs was plotted as frequency distributions for young and aged samples. Skewness was calculated in R (version 4.3.1) to evaluate the direction of distribution. A higher skewness value indicated an asymmetrical distribution biased toward smaller SG areas.

### *Drosophila* culture: Fly stocks

Canton-S and w118 wild-type *Drosophila* were maintained at 25 °C for egg laying. After eclosion, adult flies were transferred to fresh cornmeal media vials (Lakhotia & Ranganath Experiments with *Drosophila*) and subsequently moved to clean media vials every 5^th^ day to allow natural aging. Flies aged 3–5 days were used as young controls, whereas flies aged 40–50 days were designated as aged flies.

### Dissection of the *Drosophila* brain and Malpighian tubules

Control and adult *Drosophila* were anesthetized by placing them on ice. A pre-cooled Maximov cavity slide containing PBS was used to maintain the flies at low temperature during dissection.

Using fine forceps, wings and legs were removed to expose the thorax and head. The head capsule was carefully cut along the anterior edge to expose the brain, which was gently lifted with forceps or a fine needle, taking care to avoid tearing.

For abdominal dissection, a small incision was made on the ventral side with a needle tip, allowing the viscera to be gently squeezed out. The gut and attached MTs were isolated from the viscera, and MTs were further separated from the gut using fine needles. Dissected tissues were fixed in 4% formaldehyde prepared in PBS for 20 minutes at room temperature in the Maximov cavity slide.

### Immunohistochemistry of young and adult *Drosophila* tissues

Fixed brain tissues and Malpighian tubules (MTs) were washed three times with 0.3% PBST (1× PBS + 0.3% Triton X-100) for 10 minutes each. Tissues were then treated with 200 µL of 0.3% PBST three times for 10 minutes, followed by blocking in 1× PBS containing 2% BSA for 5–6 h at room temperature.

After blocking, tissues were incubated overnight at 4 °C with chicken anti-Atx2 antibody (a kind gift from Dr. Baskar Bakthavachalu, IIT Mandi, Bakthavachalu et al. 2018). The following day, primary antibody detection was performed by incubating tissues with an isotype-specific fluorescently conjugated secondary antibody (1:1000 dilution) and DAPI (1:1000 dilution) for nuclear staining for 3 h at room temperature.

Finally, tissues were washed thoroughly with 1× PBS and mounted on clean glass slides using mounting medium (10% w/v Mowiol, 25% glycerol, 2.5% DABCO in 0.1 M Tris-Cl, pH 8.5). Imaging was performed using Olympus FV3000 and Nikon ECLIPSE Ti2 microscopes. Imaging parameters were kept constant across samples to enable direct comparison of Ataxin intensity between young and aged fly brains.

### Free-floating Mice brain immunohistochemistry

Free-floating brain sectioning was performed to facilitate immunohistochemical analysis of mouse cerebellar tissue, following Tu et al. 2021. Young (2-month-old) and aged (22–24-month-old) C57BL/6J mice were transcardially perfused with 0.9% saline followed by 4% paraformaldehyde. Cerebella were extracted, post-fixed overnight at 4 °C, and cryoprotected in 30% sucrose until fully saturated.

Sagittal cerebellar sections (15 µm thickness) were obtained using a Leica CM1530 cryostat and transferred to 1× PBS. For immunostaining, sections were incubated in blocking solution containing normal donkey serum and Triton X-100, then incubated overnight at 4 °C with a primary antibody against G3BP1 (ab56574, Abcam). After extensive washing, sections were incubated with fluorophore-conjugated secondary antibodies (mouse FITC; product #715-095-150, Jackson ImmunoResearch) and co-stained with DAPI for nuclear visualization.

Stained sections were mounted in DABCO mounting medium and imaged using a Nikon AXR confocal microscope with NIS software.

### Western blot

#### Protein Isolation

Approximately 1 million cells were cultured overnight in 6-well dishes prior to trypsinization for protein isolation. Cell pellets were washed with PBS and lysed in 50–100 µL CST lysis buffer (20 mM Tris-HCl, pH 7.5; 150 mM NaCl; 1 mM Na₂EDTA; 1 mM EGTA; 1% Triton; 2.5 mM sodium pyrophosphate; 1 mM β-glycerophosphate; 1 mM Na₃VO₄; 1 µg/ml leupeptin). Lysates were centrifuged at 1300 rpm for 10 minutes at 4 °C, and the supernatant was collected in fresh tubes.

#### Protein quantification (Bradford Assay)

Total protein concentration was determined using the Bradford assay. A BSA stock solution (1 mg/ml in Milli-Q water) was serially diluted to concentrations ranging from 0.05–0.5 mg/ml to generate a standard curve. Bradford reagent was diluted 1:5 in Milli-Q water, and 200 µL of reagent was added to both standards and unknown protein samples (1:10 dilution of lysates). After 10 minutes of incubation in the dark, absorbance was measured at 545 nm using a Tecan plate reader. Protein concentrations were calculated from the BSA standard curve.

#### SDS PAGE and Protein Transfer

Equal amounts of protein (70–100 µg) were mixed with 1× loading buffer (250 mM Tris-Cl, pH 6.8; 8% SDS; 0.1% bromophenol blue; 40% v/v glycerol; 100 mM DTT) and heated at 95 °C for 10 minutes. Samples were resolved on 10% SDS-PAGE gels in 1× SDS running buffer (25 mM Tris base, 190 mM glycine, 0.1% SDS) at 100 V. Proteins were transferred to nitrocellulose membranes using the wet transfer method in 1× transfer buffer (25 mM Tris base, 190 mM glycine, 20% methanol) at 100 V for 2–3 h at room temperature.

#### Immunoblotting

Membranes were blocked with 5% BSA in 1× TBST (20 mM Tris, pH 7.5; 150 mM NaCl; 0.1% Tween-20) for 1 h at room temperature, then incubated overnight at 4 °C with primary antibodies (1:5000 dilution in 5% BSA/TBST) under gentle rocking (see antibody details in the antibody section). Membranes were washed three times with TBST (5 minutes each) and incubated with HRP-conjugated secondary antibodies (1 h, room temperature, gentle rocking). After three additional TBST washes (5 minutes each), membranes were exposed to Clarity™ ECL substrate (1:1 luminol and hydrogen peroxide) for a few seconds. Chemiluminescence was detected using a Bio-Rad Chemidoc imager.

#### Data analysis

Protein band intensities were quantified using ImageJ software. Each band was normalized to the corresponding housekeeping protein (GAPDH) within the same lane.

#### Half-Fluorescence recovery after photobleaching

HeLa G3BP1-mCherry BAC stable cells were seeded onto 22 mm diameter No. 1 circular coverslips, mounted in autoclaved stainless-steel live-cell imaging dishes, and cultured overnight at 37 °C with 5% CO₂. The following day, cells were exposed to 0.25 mM NaAsO₂ for 1 h under the same incubation conditions to induce SGs, followed by imaging at 37 °C and half-FRAP analysis.

Half-FRAP experiments were performed on a Nikon Ti2E microscope. Pre-bleach images (∼5 frames) of SG-containing cells were acquired, after which a short laser pulse (15–50% of 561 nm laser power at the source (150 mW), 50 ms duration) was applied to partially bleach SGs. Fluorescence recovery of G3BP1 was then recorded for 1 minute at low excitation intensity (15% of 561 nm LED power, exposure of 800 ms) to minimise additional bleaching. Neighbouring unbleached cells served as controls for the bleaching rate.

Pixel intensities within the bleached region of interest (ROI) were quantified using a custom MATLAB (R2023a) script. Intensities were background-corrected, normalized against a non-bleached ROI, minutes–max scaled, and subjected to curve-fitting analysis.

Recovery kinetics were modelled using a double exponential equation optimized with the Levenberg–Marquardt algorithm (Bajur et al. 2019):

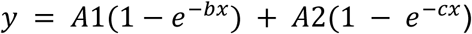

Here, recovery is represented as the sum of two fractions:

● Slow fraction: defined by parameters A1 and b, corresponding to protein diffusion from outside the granule.
● Fast fraction: defined by parameters A2 and c, corresponding to protein diffusion within the granule.
● A1 and A2 denote the amplitudes of the slow and fast fractions, respectively, while recovery times are given by the reciprocals of b and c.

### Total cellular RNA concentration determination

#### Method 1 – RNA-specific Dye staining

As one approach to quantify total RNA, we used an RNA-specific dye, SYTO® RNASelect™ green-fluorescent cell stain, to measure total cellular RNA. Young and aged HeLa Kyoto cells and human fibroblasts (∼0.05 million cells) were cultured on 12 mm-diameter No. 1 glass coverslips placed in 24-well plates. Cells were fixed with pre-chilled methanol at –20 °C for 10 minutes. Similarly, *Drosophila* brain tissues were fixed under identical conditions. Following fixation, samples were washed in PBS for 5 minutes and incubated in 40 µL RNA labeling solution containing 5 µM SYTO® RNASelect™ green fluorescent cell stain (prepared from a 5 mM stock) and 1 µg/ml DAPI in PBS for 20 minutes at room temperature. The labeling solution was freshly prepared and used immediately to avoid dye precipitation and debris formation on coverslips. After staining, cells and tissues were washed three times in PBS, mounted with Fluoromount-G™ mounting medium, and sealed with transparent nail polish. Imaging of RNA-labeled cells was performed immediately using Nikon ECLIPSE Ti2 and *Drosophila* brain samples in Olympus FV3000 microscopes.

#### Method 2-Metabolic labeling of total RNA

As a second approach, RNA was metabolically labeled in vivo using EU, a uridine analogue incorporated into RNA during transcription. Young and aged HeLa Kyoto cells, human fibroblasts, and mouse fibroblasts (∼ 0.05 million cells) were cultured on 12 mm diameter No. 1 glass coverslips placed in 24-well plates. Cells were incubated with 1 mM EU (prepared from a 100 mM stock in Milli-Q water) in culture medium at 37 °C with 5% CO₂ for 24 h to label total RNA.

Following incubation, cells were washed three times with PBS and fixed with 4% formaldehyde for 10 minutes at room temperature. Permeabilization was performed using 0.25% Triton X-100 in PBS for 10 minutes, after which cells were rinsed with 300 mM glycine for 5 minutes and blocked with 2% BSA in PBS for 2 h. After blocking, cells were washed three times with 100 mM HEPES. EU-labeled RNA was detected via Alkyne–Azide click chemistry (Salic and Mitchison, 2008) using Alexa 555 fluorescent azide. A 200 µL click reaction mixture containing 100 mM HEPES (pH 7.25), 2 µM Alexa 555 azide, 1 mM CuSO₄, 0.5 mM THPTA, 5 mM ammonium guanidine hydrochloride, and 5 mM sodium ascorbate was added to the cells and incubated in the dark for 2 h.

After reaction, coverslips were washed with PBS, mounted in Fluoromount-G™ mounting medium, and sealed with transparent nail polish. Imaging of EU-labeled RNA was performed using Nikon ECLIPSE Ti2 and Leica DMI6000b microscopes.

#### Method 3 - RNA isolation and quantification

A third approach to quantify RNA was to isolate RNA from an equal number of cells and quantify it following gel electrophoresis. Control and senescent HeLa Kyoto cells (∼ 1 million cells) were trypsinized, and cell pellets were washed with PBS. RNA was extracted using TRIZOL reagent: 1 ml TRIZOL was added to each pellet and incubated at room temperature for 5 minutes. Subsequently, 0.2 ml chloroform was added, and samples were centrifuged at 12,000 g for 15 minutes at 4 °C. Phase separation yielded three layers: the upper aqueous phase (RNA), the interphase (DNA), and the lower organic phase. The RNA-containing aqueous layer was carefully transferred to a fresh tube, avoiding contamination from the middle layer. RNA was precipitated by adding 0.5–1 ml of isopropanol, mixing thoroughly, and incubating for 10 minutes at room temperature. Samples were centrifuged at 12,000 g for 10 minutes at 4 °C, and the supernatant was discarded. Pellets were washed with 75% cold ethanol by centrifugation at 7,500 g for 5 minutes at 4 °C. After discarding the supernatant, pellets were air-dried for several minutes and resuspended in 25 µL RNase-free water. For electrophoresis, 1–2 µL of RNA was loaded onto the gel. RNA bands were visualized and quantified using ImageJ software.

#### RNA transcription rates

Approximately 0.05 million cells were seeded onto round No. 1 glass coverslips (12 mm diameter) placed in a 24-well plate. Cells were incubated with 1 mM EU, prepared in culture medium from a 100 mM stock solution (dissolved in Milli-Q water), at 37 °C with 5% CO₂ for 30 minutes to label nascent RNA transcripts. The labeled RNA was subsequently detected using an alkyne–azide click chemistry ligation reaction, as described above.

#### RNA degradation rates

Approximately 0.05 million cells were plated onto round No. 1 glass coverslips (12 mm diameter) placed in a 24-well plate. To assess RNA degradation rates, newly synthesized RNA was metabolically labeled as described above. Following 30 minutes of EU incorporation, the EU-containing medium was replaced with medium supplemented with 10 µg/mL Actinomycin D to inhibit transcription and enable measurement of RNA decay. Cells were treated with Actinomycin D for 1 or 2 hours at 37 °C in 5% CO₂, then fixed with 4% formaldehyde for 10 minutes at room temperature. After fixation, samples were processed to detect the labeled RNA using an alkyne–azide click chemistry ligation reaction, as described above.

#### Quantitative PCR

Senescence validation was performed by assessing the expression of cell cycle arrest markers (P21, P16) and pro-inflammatory cytokines (IL-6, IL-8) using real-time quantitative PCR (see primers section). Total RNA (500 ng) was reverse-transcribed into cDNA using the PrimeScript™ cDNA Synthesis Kit. The resulting cDNA was subjected to quantitative PCR for gene expression analysis. Reactions were carried out on a Bio-Rad CFX Opus 384 Real-Time PCR system using either TB Green® Premix Ex Taq™ (Tli RNase H Plus) or Luna® Universal qPCR Master Mix in a 10 µl reaction volume. GAPDH expressions were used as an internal control for normalization. Data and melting curve analyses were performed using Bio-Rad CFX Manager software.

#### NHS Ester protein labeling

To assess protein density within SGs during senescence, amino acids were labeled, following the protocol adapted from (Sheard et al. 2023). Control and senescent HeLa Kyoto cells (∼50,000 cells) were cultured on 12 mm-diameter No. 1 glass coverslips in 24-well plates. Cells were stressed for 1 hour with 0.25 mM NaAsO₂, then washed three times with PBS and fixed with 4% formaldehyde for 10 minutes at 37 °C and 5% CO₂. Fixed cells were incubated with 1 mM NHS-Fluorescein (5/6-carboxyfluorescein succinimidyl ester) prepared in DMSO for 30 minutes at 37 °C and 5% CO₂. Staining was quenched with 1 M Tris-Cl (pH 8.0) for 15 minutes at 37 °C. Cells were then permeabilized with 0.25% Triton X-100 for 10 minutes at room temperature, followed by blocking in 2% BSA in PBS for 1–2 hours at room temperature. Subsequently, cells were incubated with primary antibodies diluted in 2% BSA for 16 hours at 4 °C. After three PBS washes, coverslips were incubated with Alexa Fluor™-conjugated secondary antibodies (1:1000 dilution in 2% BSA/PBS) for 1 hour at room temperature. Nuclear staining was performed with 1 µg/mL DAPI in PBS for 10 minutes at room temperature. Coverslips were washed three times with PBS, mounted in Fluoromount-G™ medium, and sealed with transparent nail polish. Imaging was performed using a Nikon ECLIPSE Ti2 microscope equipped with an ORCA®-Fusion BT camera.

### G3BP1 protein purification

#### Cloning MBP-G3BP1-GFP-His

For bacterial expression of G3BP1, we engineered a fusion construct in which the MBP sequence was fused to the N-terminus of G3BP1, while the EGFP sequence and a His tag were fused to the C-terminus. The G3BP1 coding sequence was flanked by EcoRI and AscI restriction sites.

The construct design was based on elements from the plasmid TH0780-POEM1-MBP-PreScission-FUS-TEV-EGFP-PreScission-His (a kind gift from Prof. Anthony Hyman’s lab, MPI-CBG, Dresden; Patel et al. 2015). From this plasmid, we adopted the N-terminal MBP tag, the C-terminal EGFP-His tag, and the PreScission protease and TEV protease cleavage sites. The resulting fusion construct, MBP-PreScission-G3BP1-TEV-EGFP-PreScission-His, was subsequently cloned into the pET30a vector backbone (a kind gift from Prof. Simon Alberti, TU Dresden).

In parallel, the PreScission protease sequence was amplified from a vector encoding the protease fused to a GST tag (a kind gift from Prof. Mahavir Singh’s lab, Indian Institute of Science, Bengaluru, India). This amplified fragment was then inserted into the pET-Duet 1 vector (a kind gift from Prof. B. Gopal’s lab, Indian Institute of Science, Bengaluru) using BamHI and HindIII restriction sites.

#### G3BP1 protein purification

The pET30a(+) vector containing the MBP-G3BP1-GFP-His construct was transformed into *E. coli* Rosetta(DE3)pLysS cells (a kind gift from Prof. B. Gopal’s lab, Indian Institute of Science, Bengaluru, India) (Millipore) for fusion G3BP1 protein expression. Cells were cultured in Luria–Bertani (LB) broth to an OD₆₀₀ of 0.6 and induced with 0.5 mM IPTG at 16 °C overnight. Following induction, cells were pelleted, resuspended, and lysed in buffer containing 50 mM HEPES (pH 7.5), 300 mM NaCl, 5% glycerol, 20 mM imidazole, 1 mM β-mercaptoethanol, and complete protease inhibitor cocktail (Roche). Lysates were clarified by centrifugation at 11000 RCF for 45 minutes at 4 °C.

The supernatant was incubated with Ni-NTA beads (G-Biosciences, 786-1547) for 1 hour, and bound protein was eluted using a gradient of imidazole (up to 500 mM). Elution fractions were analyzed by SDS-PAGE, pooled, and dialyzed against buffer containing 50 mM HEPES (pH 7.5), 300 mM NaCl, 5% glycerol, and 1 mM DTT. The purified protein was concentrated, flash-frozen in liquid nitrogen, and stored at −80 °C.

PreScission protease was similarly expressed and purified from Rosetta (DE3) pLysS cells using Ni-NTA affinity chromatography. For this purification, cells were lysed in buffer containing 50 mM Tris-HCl (pH 7.5), 500 mM NaCl, and 10% glycerol. The purified protease was stored at −80 °C in buffer containing 50 mM Tris (pH 7.0), 150 mM NaCl, and 5% glycerol. Protein concentrations were determined using the Bradford assay, as described above.

### In vitro lysate experiments

#### Cell lysate preparation

HeLa Kyoto cells (2-3 million cells) were trypsinized and pelleted at 200 RCF for 4 minutes, followed by a PBS wash. After removal of PBS, the cell pellets were stored at −80 °C until further use. Frozen pellets were later thawed at room temperature for 5 minutes. Cells were lysed in buffer containing 50 mM Tris (pH 7.0), 0.5% NP-40, and 2% recombinant RNase inhibitor. Lysis was performed by pipette mixing until a homogenate was obtained, followed by incubation at room temperature for 5 minutes. Lysates were clarified by centrifugation at 21,000 g for 5 minutes, and the supernatant was transferred to a fresh centrifuge tube for use in vitro experiments. This protocol was adapted from Freibaum et al. 2021.

#### In vitro Stress granule reconstitution

G3BP1 condensates were induced by adding 5% PEG to a reaction mixture containing preScission protease (in-house purified, used to cleave MBP and His tags from the MBP-G3BP1-GFP-His construct), 3.5 µM purified G3BP and 100 ng of total RNA isolated from HeLa Kyoto cells. SG-like condensates were induced by adding cellular lysates to the above reaction mixture.

A 10 µl reaction volume was immediately dispensed into 384-well Greiner Bio-One plates (#781906). Droplets were allowed to settle for 30 minutes before imaging on a Nikon ECLIPSE Ti2 fluorescence microscope with a 60× oil-immersion objective and 488 nm excitation. G3BP1 and SG-like condensate intensity and enrichment were quantified using custom MATLAB (R2023a) scripts.

#### FRAP to study In vitro stress granule dynamics

Half-FRAP experiments of in vitro G3BP1 and SG-like condensates were performed on a Nikon Ti2E microscope. Pre-bleach images (∼5 frames) of condensates were acquired, after which a short laser pulse (100% of 488 nm laser power at the source, 50 ms duration) was applied to partially bleach the condensates. Fluorescence recovery of G3BP1 was then recorded for 2 minutes at low excitation intensity (15% of 488 nm LED power) to minimize additional bleaching. Neighboring unbleached condensates served as controls for the bleaching rate.

Pixel intensities within the bleached region of interest (ROI) were quantified using a custom MATLAB R(R2023a) script. Intensities were background-corrected, normalized against a non-bleached ROI, minutes–max scaled, and subjected to curve-fitting analysis. The recovery kinetics were further calculated using the method mentioned above.

#### Quantification and statistical analysis

The comparison of mean intensities was carried out using the Mann-Whitney U test and the Student’s t-test. p<0.05 was considered significant. Data are presented as mean ± SD or mean ± SEM as indicated.

#### Animal Research Ethics

This protocol was approved by the Ethics Committee on the Use of Animals of the Committee for the Control and Supervision of Experiments on Animals (CCSEA), under the approved protocol issued by the Institutional Animal Ethics Committee of Indian Institute of Science, Bangalore (CAF/ETHICS/983/2023) and Indian Institute of Technology, Kanpur (IITK/IAEC/2024/1228).

